# Lineage plasticity of the integrated stress response is a hallmark of cancer evolution

**DOI:** 10.1101/2025.02.10.637516

**Authors:** Shiqi Diao, Jia Yi Zou, Shuo Wang, Nour Ghaddar, Jason E. Chan, Hyungdong Kim, Nicolas Poulain, Constantinos Koumenis, Maria Hatzoglou, Peter Walter, Nahum Sonenberg, John Le Quesne, Tuomas Tammela, Antonis E. Koromilas

## Abstract

The link between the “stress phenotype”-a well-established hallmark of cancer-and its role in tumor progression and intratumor heterogeneity remains poorly defined. The integrated stress response (ISR) is a key adaptive pathway that enables tumor survival under oncogenic stress. While ISR has been implicated in promoting tumor growth, its precise role in driving tumor evolution and heterogeneity has not been elucidated. In this study, using a genetically engineered mouse models, we demonstrate that ISR activation—indicated by elevated levels of phosphorylated eIF2 (p-eIF2) and ATF4—is essential for the emergence of dedifferentiated, therapy-resistant cell states. ISR, through the coordinated actions of ATF4 and MYC, facilitates the development of tumor cell populations characterized by high plasticity, stemness, and an epithelial-mesenchymal transition (EMT)-prone phenotype. This process is driven by ISR-mediated expression of genes that maintain mitochondrial integrity and function, critical for sustaining tumor progression. Importantly, genetic, or pharmacological inhibition of the p-eIF2–ATF4 signaling axis leads to mitochondrial dysfunction and significantly impairs tumor growth in mouse models of lung adenocarcinoma (LUAD). Moreover, ISR-driven dedifferentiation is associated with poor prognosis and therapy resistance in advanced human LUAD, underscoring ISR inhibition as a promising therapeutic strategy to disrupt tumor evolution and counteract disease progression.

## INTRODUCTION

Cancer is a complex genetic disease driven by gain-of-function mutations, amplification, or overexpression of oncogenes, alongside loss-of-function mutations, deletions, or epigenetic silencing of tumor suppressor genes ^1^. A hallmark of cancer is the “stress phenotype,” arising from increased DNA damage, proteotoxicity, and metabolic stress due to oncogenic mutations or factors in the tumor microenvironment (TME) ^2^. To proliferate, tumor cells rely on internal pathways that promote survival and adaptation under these stressful conditions.

The integrated stress response (ISR) is a vital cellular mechanism that enables cells to adapt to various stressors, such as nutrient deprivation, hypoxia, and misfolded protein accumulation ^3^. By modulating protein synthesis, the ISR supports cell survival under manageable stress and initiates processes to eliminate irreparably damaged cells under severe stress ^3^. Key stress-sensing kinases, including PERK, GCN2, PKR, and HRI, activate the ISR by phosphorylating the α subunit of eIF2 at serine 52 (p-eIF2α) ^4,5^. This phosphorylation decreases overall protein synthesis while selectively enhancing the translation of stress-related proteins like ATF4, a transcription factor involved in cellular adaptation and survival ^6^.

ISR protects normal cells from the damaging effects of stress, thereby preventing tumorigenesis ^3,4,7^. However, cancer cells can exploit the ISR to enhance their survival and proliferation within the often-hostile TME, characterized by hypoxia, nutrient scarcity, and oxidative stress ^4,8^. ISR activation and the resulting translational reprogramming enhance the efficiency of oncogenic mRNA translation and promote tumor cell survival ^9–11^, establishing translational control as a critical therapeutic target ^12^. Cancer cells leverage the ISR to sustain growth ^4^, presenting a promising avenue for therapeutic intervention ^11,13,14^.

The translational and transcriptional programs activated by the ISR are intricately connected with other cellular stress responses, such as the unfolded protein response (UPR) ^15^ and oxidative stress responses^16^, to ensure a coordinated and comprehensive reaction to various types of cellular and oncogenic forms of stress.

While elevated ISR activity has been observed in tumor cells exhibiting stemness ^17^, its role in driving tumor progression and dedifferentiation associated with therapy resistance remains unclear. To address this, we focused on lung adenocarcinoma (LUAD), which represents 35–40% of all lung cancers and is often diagnosed at advanced stages, with a median survival of just over 18 months ^18,19^. The limited effectiveness of current therapies against LUAD underscores the urgent need for more effective treatment strategies ^20^.

Previous work using single-cell transcriptomics in mouse LUAD models has identified highly plastic cell states that promote tumor progression, intratumor heterogeneity, and therapy resistance through mechanisms like lineage switching and epithelial-mesenchymal (EMT) transition ^21–25^. While ISR’s role in lineage transitions during tumor progression has been unclear, our investigation using lineage tracing and single-cell transcriptomics reveals ISR’s critical role in driving LUAD expansion into cell lineages characterized by high plasticity and stemness. This process is mediated through the activity of transcription factors ATF4 and MYC.

In this model, which is significantly impacted by both genetic and pharmacological inhibition of key components of ISR, like p-eIF2α and ATF4, we discovered a new cluster of tumor cells that represent a developmental dead-end owing to ISR suppression. This cluster is characterized by disrupted mitochondrial function, which is essential for meeting the energy demands and survival of cancer cells. These findings demonstrate that the ISR not only promotes tumor plasticity and stemness but also plays a key role in the developmental trajectory of LUAD. Interestingly, pharmacological inhibition of ISR by ISRIB, a small molecule with proven efficacy in treating cognitive disorders ^3^, effectively stalls the evolutionary process of tumors. This positions ISRIB as a promising therapeutic candidate for strategies aimed at halting tumor progression and metabolic adaptation.

## RESULTS

### Loss of p-eIF2α disrupts LUAD tumor evolution in KP mice

The KP model is a genetically engineered mouse model of lung cancer that mimics human LUAD, characterized by the Cre-loxP-mediated expression of oncogenic *KRAS G12D* and loss of the TP53 tumor suppressor gene ^26^. In this study, the KP model was modified by introducing a conditional homozygous S52A mutation in the *Eif2s1* allele (fTg/0; eIF2α^A/A^) ^11^, which prevents phosphorylation of serine 52 and inhibits ISR activation. Mice with a wild-type *Eif2s1* allele (fTg/0; eIF2α^S/S^) were used as controls.

Lung tumors were induced using intratracheal intubation of CRE-expressing lentiviruses driven by the carbonic anhydrase 2 promoter, targeting alveolar type 1 and 2 lung cells (Fig. 1a) ^11^. These lentiviruses also expressed *Trp53* shRNA to accelerate tumor formation ^11^. GFP expression was specifically enabled in lung tumors for visualization and sorting via flow cytometry (Fig. 1a) ^11^. Noninvasive ultrasound imaging revealed that loss of p-eIF2α significantly reduced lung tumor formation and growth in live mice, consistent with prior findings (Fig. 1b) ^11^.

**Fig. 1.**
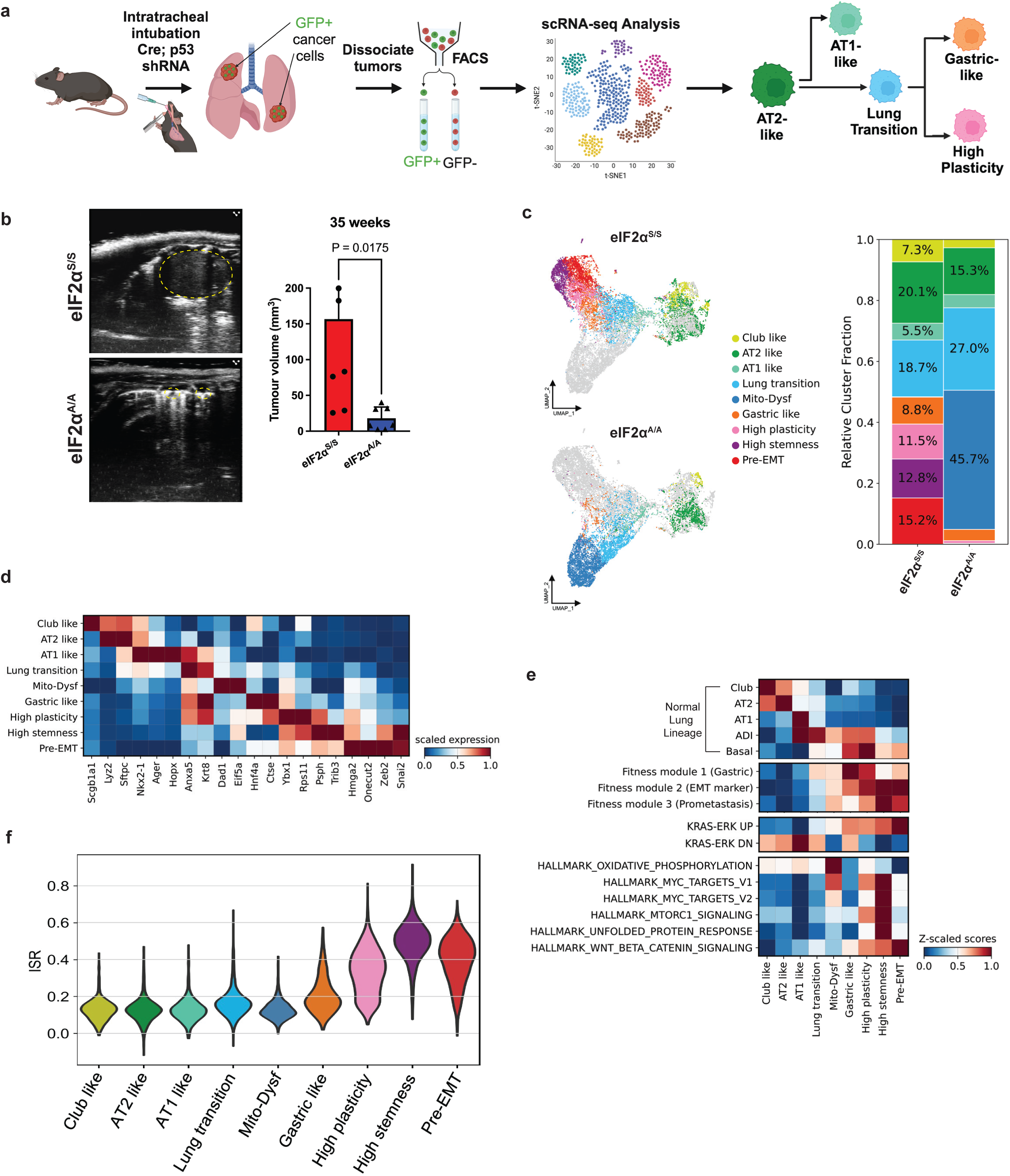
The loss of p-eIF2α disrupts the progression of LUAD tumors to high-fitness lineages, leading to decreased tumor heterogeneity in KP mice. (**a**) Schematic overview of the experimental design and analysis workflow. The schematic was created using BioRender (https://www.biorender.com). (**b**) Representative ultrasound images show mouse lung tumors detected in the septum, located peripherally in contact with the pleura, 35 weeks after CRE-lentivirus intubation in eIF2α ^S/S^ (n = 6) and eIF2α ^A/A^ (n = 7) mice. Tumor is outlined with a yellow intermittent line. Data were analyzed using the Mann-Whitney U Test and are presented as mean ± SD. (**c**) Uniform manifold approximation and projection (UMAP) of tumor cells colored by annotated cell types in respective genotypes with fractions of each cluster in different tumor type (n= 17,919 cells). (**d**) Heatmap illustrating the average expression levels (scaled) of selected marker genes across identified clusters. This visualization highlights distinct marker gene enrichment patterns, aiding in the classification of cellular heterogeneity within the dataset. (**e**) The signature scores of selected gene sets and hallmark pathways were calculated across the annotated cell types. (**f**) A comparison of ISR scores between annotated cell types from the merged eIF2α^S/S^ and eIF2α^A/A^ tumor datasets reveals differences in ISR activity across the tumor cluster types.

Previous studies on tumor evolution in KP mice have shown that alveolar type 2 (AT2) cells transformed by mutant KRAS undergo lineage transitions and phenotypic changes, eventually acquiring states of high plasticity and stemness, which are key drivers of tumor heterogeneity^22–24^.

To investigate the role of ISR, particularly p-eIF2α, in LUAD evolution, GFP+/CD45-lung tumor cells of our KP model were isolated via flow cytometry and subjected to single-cell (sc) RNA-seq analysis (Fig. 1a). To ensure sufficient representation of cells with impaired p-eIF2α, we selected tumors from mice that had been grown for 35 weeks, as tumors with impaired p-eIF2α exhibit significant growth inhibition (Fig. 1b). Our analysis, which consisted of unsupervised clustering ^27^ and automatic cell type annotation ^28^, identified nine clusters with unique expression patterns (Fig. 1c,d and Supplementary Table 1). We analyzed each cell subgroup by identifying differentially expressed genes and comparing their profiles to published scRNA-seq datasets from mouse cell atlases ^21–24^.

Enrichment analysis revealed that some tumor cell populations—such as club-like, AT2-like, AT1-like, lung transition, and gastric-like clusters—corresponded closely to normal lung or gastric cell marker gene sets, indicating high differentiation characteristics (Fig. 1c,d). Other tumor cell populations, including clusters characterized by high plasticity, stemness, and pre-EMT traits, did not align with normal lineage markers. Instead, these populations displayed traits resembling placental or embryonic cells, reflecting significant lineage distortion and an undifferentiated state (Extended Data Fig. 1a and Supplementary Table 1). Six of the identified clusters—AT2-like, AT1-like, club-like, gastric-like, high plasticity, and pre-EMT—correspond to cell states previously described in fully formed KP LUAD tumors^23^ (Fig. 1d, Extended Data Fig. 1b and Supplementary Table 1). Notably, the club-like cells differ from primitive club cells, representing a subset of AT2-like cells marked by the expression of *Scgb1a1* and AT2-like cell marker *Sftpc* (Fig. 1d, Extended Data Fig. 1b and Supplementary Table 1).

We identified an early cluster, termed the lung transition cluster, which exhibits characteristics of AT2- and AT1-like clusters (Fig. 1c). This transition state is marked by the co-expression of AT1 markers like *Hopx* and *Ager* alongside AT2 markers like *Sftpc* and *Lyz2*, as well as elevated levels of the *Krt8,* a marker of AT2-AT1 intermediate cells ^29^ (Fig. 1d and Supplementary Table 1). This state resembles the intermediate epithelial cell population between AT2 and AT1 cells during alveolar development in mice impaired for *Ndufs2* ^29^.

Former studies of normal lung epithelial cells and mouse lung tumors identified a type of undifferentiated cell with characteristics of high stemness ^25^. Our scRNA-seq data similarly identified a high stemness cell cluster, characterized by upregulated unfolded protein response (UPR), mTOR signaling, and MYC-dependent genes (Fig. 1e). This late undifferentiated state does not correspond to any specific lung epithelial lineage, like AT2 and AT1 cells (Fig. 1e, top panel). Instead, it is characterized by relatively moderate activation of basal stem cell programs and elevated MYC target gene expression (Fig. 1e), features commonly observed in normal tissue stem cells ^30^. We refer to it as the high stemness cluster. Notably, this cluster exhibits a high transcriptomic similarity to the pre-EMT cluster, as evidenced by a shorter Bhattacharyya distance, which measures the overlap in the probabilistic distributions of the clusters ^31^, compared to other clusters (Extended Data Fig. 1c). However, the pre-EMT cluster expresses more neuronal reprogramming genes, such as *Hmga2* and *Onecut2* ^32,33^, as well as EMT markers like *Zeb2*^34^ (Fig. 1d and Supplementary Table 1).

KP tumors with impaired ISR due to homozygous S52A knock-in mutation of *Eif2s1*—referred to as eIF2α^A/A^ tumors—contained the AT1-like, AT2-like, and lung transition clusters, along with small populations of the gastric-like and club-like clusters. Additionally, these cells exhibited the presence of a new cluster characterized by the dysregulation of nuclear-encoded mitochondrial genes (Fig. 1c). This late cluster, designated as the Mito-Dysf cluster, exhibited increased expression of lung lineage-defining transcription factors *Nkx2-1*^35,36^ and AT2 markers like *Sftpc* and *Lyz2* compared to high stemness and pre-EMT cells (Fig. 1 and Supplementary Table 1). In eIF2α^S/S^ tumors that can phosphorylate eIF2α, a switch from lung lineage to primitive gut or developmental lineage factors is often characteristic of late tumors (Fig. 1d and Extended Data Fig. 1a). However, Mito-Dysf cluster also displayed a significantly reduced expression of the developmental master regulators *Hnf4a*, associated with the primitive gut ^37^, and *Hmga2,* which is linked to the primitive gut and developing lung ^38^. Using the Weissman’s fitness module 1-3 ^24^, these cells did not align with any established tumor evolutionary fates, suggesting they may represent a failed cellular reprogramming process (Fig. 1e, second panel from the top).

We applied an ISR transcriptional signature to assess ISR activity signature score across various KP lung tumor clusters ^29^. This gene set consists of upregulated ISR target genes identified through an unbiased, tissue-independent approach *in vivo*, under conditions without the influence of exogenously applied pleiotropic stressors ^29,39^. The findings revealed low ISR signature score in the Mito-Dysf, AT2-like, AT1-like, and gastric-like clusters, intermediate ISR signature score in the lung transition cluster, and elevated ISR signature score in the high plasticity and pre-EMT clusters (Fig. 1f). Notably, the highest ISR signature score was observed in the high stemness cluster (Fig. 1f).

### ISR-upregulated genes drive lineage transition in LUAD tumors

To model the lineage transition of tumor cells, we utilized CellRank for single-cell fate mapping^40,41^. We identified five stable clusters representing AT2-like cells as the initial state and predicted four distinct terminal differentiation clusters: AT1-like, gastric-like, pre-EMT, and Mito-Dysf groups (Fig. 2a,b). Fate probability analysis showed that AT2-like cells contributed uniformly to all terminal cell fates (Fig. 2c,d). In early stage of KP tumor development, there is no difference in AT2-like cells between tumors with an intact (eIF2α^S/S^) and impaired p-eIF2α (eIF2α^A/A^) (Fig. 2d). These results suggest that the early stages of lineage trajectory of KP lung tumors do not require the ISR pathway.

**Fig. 2.**
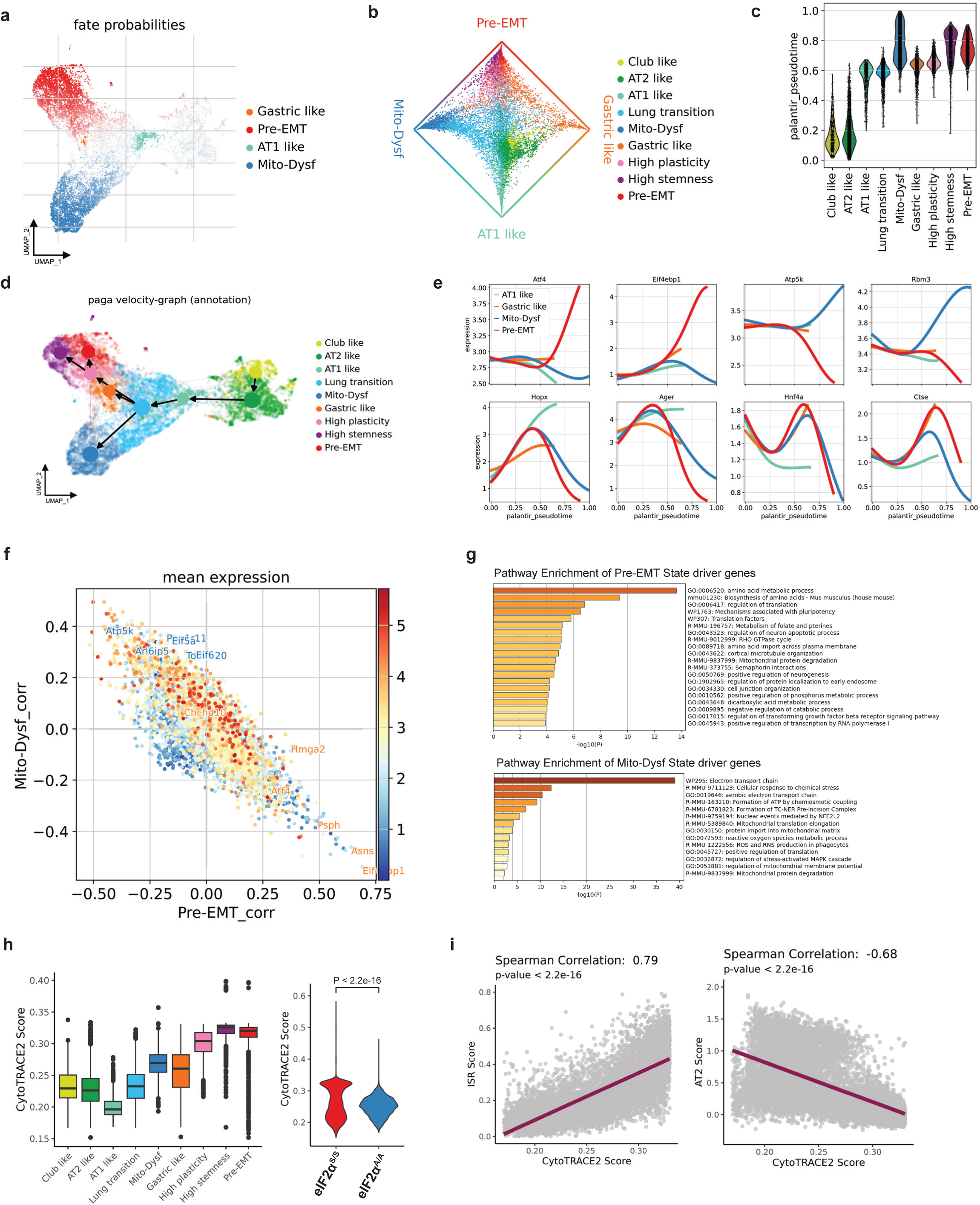
ISR signaling drives LUAD progression and acts as the primary determinant of the high-fitness lineage trajectory. (**a**) CellRank-computed macrostates highlight the most confidently assigned cells for each state. Names and colors were assigned to these macrostates based on their overlap with the clusters from Fig. 1c, with AT2-like cells representing the initial cell states. (**b**) A circular projection of cells based on their fate probabilities toward the macrostates shown in (a), with colors representing cluster annotations from Fig. 1c. Macrostates are positioned around the edge of the plot, and each cell is placed within the plot according to its likelihood of progressing toward any of the terminal states. (**c**) Pseudotime scores and predicted ordering of the 7 clusters. (**d**) The PAGA plot illustrates the directional development of tumor cell types based on RNA velocity. The velocity field is projected onto the UMAP plot, with PAGA aggregating these transitions at the cluster level. Arrows indicate the local average velocity, evaluated across a regular grid, to depict the flow of cellular transitions. (**e**) Smoothed gene expression trends along pseudotime, with each colored trend weighted by CellRank fate probabilities for the indicated lineages. The expression trend for each gene is displayed across trajectories leading to the corresponding terminal populations. (**f**) Genes that display a strong positive correlation with the pre-EMT fate are strongly negatively correlated with the Mito-Dysf fate, and vice versa, highlighting an inverse relationship between these two cellular trajectories. (g) Functional annotation of the representative top genes reveals distinct roles for pre-EMT-driven genes and Mito-Dysf-driven genes. (h) Cytotrace-based stemness scores were calculated to quantify the stemness properties across various cell populations or genotypes. The left panel illustrates the distribution of stemness scores for each identified cluster, while the right panel compares these scores across different tumor genotypes. (i) Correlation analyses between Cytotrace stemness scores, ISR activity, and AT2 signatures. Spearman’s correlation test was used to calculate p-values.

Along the trajectory, a unique lung transition cluster emerges, with entropy values starting to decrease (Extended Data Fig. 2a). Analysis of this cluster indicated that it represents a focal point for transition to the rest of the clusters (Extended Data Fig. 2b-d). The lung transition cluster is found in tumors with intact as well as impaired p-eIF2α (Extended Data Fig. 2c). However, the lung transition tumor cells with intact p-eIF2α (eIF2α^S/S^) exhibit a gradually increasing expression of ISR-dependent genes like *Asns, Eif4ebp1* (Extended Data Fig. 2d) and increased expression of genes, like *Apobec3*, and pathways with roles in chromatin remodelling (Extended Data Fig. 2d and Supplementary Table 1). On the other hand, lung transition cluster cells with loss of p-eIF2α (eIF2α^A/A^) display increased expression of genes like *Rbm3, Uba52* and *Eif5a*, with roles in mitochondrial biogenesis and function (Extended Data Fig. 2d and Supplementary Table 1) ^42–44^.

To explore gene expression dynamics along the differentiation trajectories of the four terminal clusters, we measured the dynamics of genes in the pseudotime along the differentiation trajectories. We found that genes like *Hmga2*, *Onecut2,* with high relevance to KP lung tumor evolution ^23^, were significantly upregulated in clusters of the pre-EMT trajectory (Extended Data Fig. 2e). Known ISR-upregulated genes, such as *Atf4* and *Eif4ebp1*, also increased along the pseudotime dynamics reaching high expression in the terminal pre-EMT state (Fig. 2e). These data suggest that tumor evolution and the ISR pathway are closely linked to the acquisition of pre-EMT state. Among the key driver genes associated with the pre-EMT state, the ISR-dependent gene *Eif4ebp1* emerged as a top candidate (Fig. 2f and Supplementary Table 2). The top driver gene for the AT1-like cell fate, *Hopx*, corresponds to the normal AT1 cell lineage factor. Similarly, for the gastric-like state, *Hnf4a* serves as the key lineage factor (Fig. 2e).

Conversely, genes with mitochondrial functions such as *Rbm3*, *Atp5k*, *Eif5a* and *Tomm20* were increased during the transition to Mito-Dysf state for eIF2α^A/A^ tumors devoid of p-eIF2α (Fig. 2e and Extended Data Fig. 2e). The pre-EMT and Mito-Dysf states were found to be mutually exclusive, as genes positively correlated with one state showed a negative correlation with the other (Fig. 2f). AT1-like driver genes also have negative correlation with pre-EMT driver genes (Extended Data Fig. 2f), consistent with the characteristics of pre-EMT dedifferentiation (Extended Data Fig. 1a). AT1-like and gastric-like fate had a weak correlation with the Mito-Dysf state (Extended Data Fig. 2f). Pathway enrichment revealed that top genes of the pre-EMT state are linked to mRNA translation, amino acid biosynthesis, cytoskeleton organization and cellular response to stress (Fig. 2g). Conversely, top genes of Mito-Dysf state are linked to functions like oxidative phosphorylation, ATP synthesis, mitochondrial protein synthesis, and mitochondrial protein degradation (Fig. 2g).

Next, we addressed the connection between ISR and the developmental potential hierarchy of lung tumors. We applied CytoTRACE to predict the differentiation state of each cluster from the scRNA-seq data ^45,46^. CytoTRACE analysis showed that the “stemness” score in the high stemness and pre-EMT clusters was higher than in the other clusters suggesting the association of these cells with potential stem cells (Fig. 2h, left panel). Despite being at the trajectory’s end (Fig. 2a,d), the Mito-Dysf cluster in eIF2α^A/A^ tumors consisted of cells of moderate differentiation that did not acquire a stemness phenotype (Fig. 2h). The ISR signature score is positively correlated with the stemness/dedifferentiation score (Fig. 2i) and the MYC signature score ((Extended Data Fig. 2g), indicating that MYC is likely to promote the ISR-mediated evolution of lung cancer tumors.

Using the CellRank program we identified a transcriptional signature of 160 genes driving the transition from AT2-like to pre-EMT cells, with correlation coefficients greater than 0.3 (Supplementary Table 2). Among these 160 genes, 50 genes were ISR-dependent, including *Eif4ebp1* and *Psph*, which play roles in mRNA translation and cellular metabolism, respectively (Extended Data Fig. 3a-c and Supplementary Table 2). These finding suggest a key role of ISR in driving the transition of AT2-like to pre-EMT cells. Immunohistochemical (IHC) analysis showed that 4EBP1 levels were higher in eIF2α^S/S^ than eIF2α^A/A^ tumors (Extended Data Fig. 3d). This pattern aligned with that of ONECUT2, a marker of high plasticity, stemness, and pre-EMT clusters, in these tumors (Extended Data Fig. 3d). Our findings implicate ISR in pathways facilitating the transition from AT2-like to pre-EMT cells, ultimately supporting lung tumor progression.

### Partial ISR disruption is enough to impede LUAD tumor progression

We next determined lineage transition in KP lung tumors expressing a heterozygous *Eif2s1 S52A* knock in allele (eIF2α^S/A^). Ultrasound imaging of the lungs of live mice showed that eIF2α^S/A^ tumors displayed an impaired growth compared to eIF2α^S/S^ tumors (Fig. 3a). Sc-RNA seq analyses revealed that eIF2α^S/A^ tumors contained all the clusters found in eIF2α^S/S^ tumors but lacked the Mito-Dysf cluster present in eIF2α^A/A^ tumors (Fig. 3b). Nevertheless, the high plasticity and pre-EMT clusters were substantially reduced in eIF2α^S/A^ compared to eIF2α^S/S^ tumors (Fig. 3b). Conversely, the AT2-like and gastric-like clusters were increased in eIF2α^S/A^ compared to eIF2α^S/S^ tumors, indicating that partial loss of p-eIF2α can block the transition of tumors from the differentiated AT2-like state to the fully undifferentiated pre-EMT cluster (Fig. 3b).

**Fig. 3.**
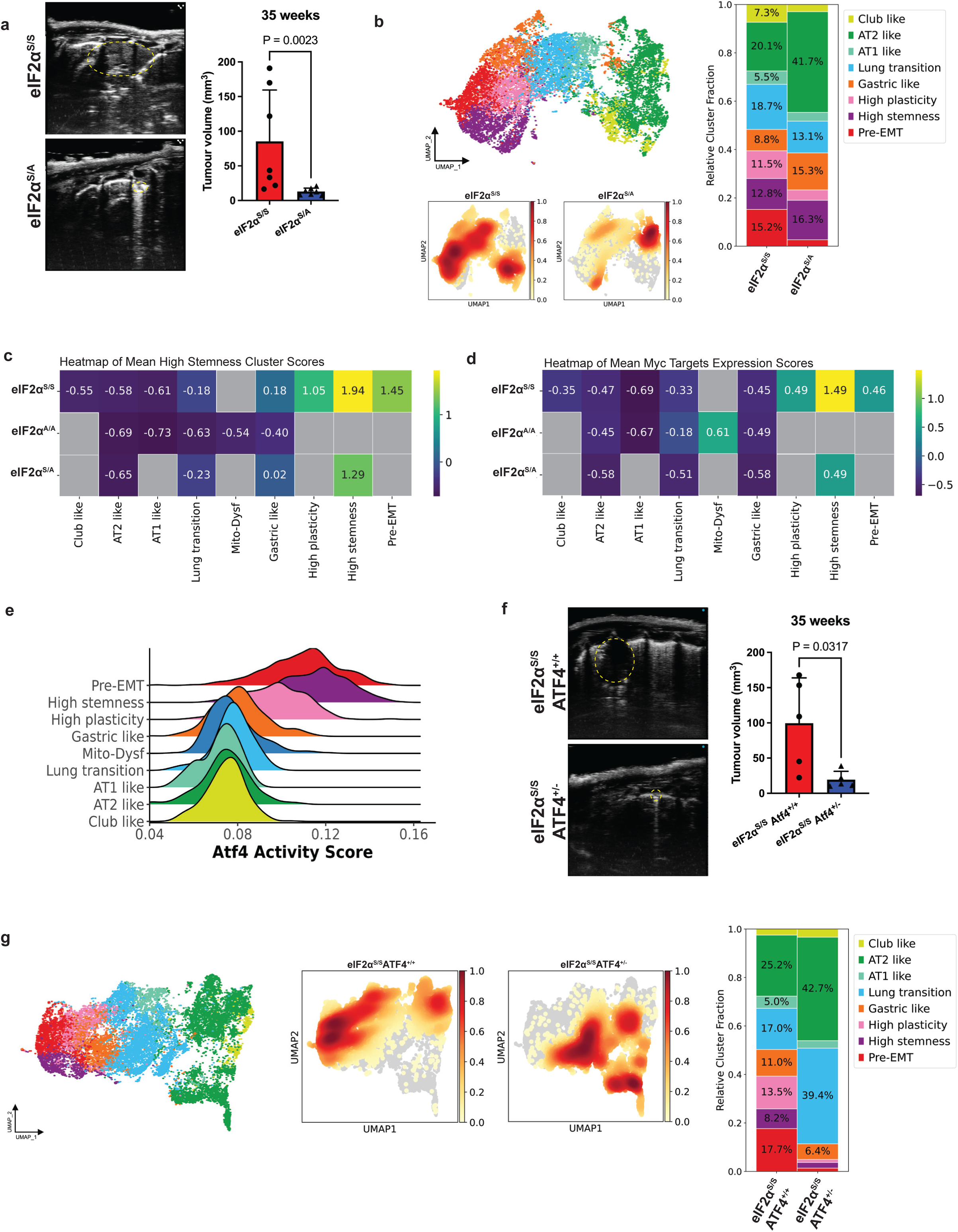
Disrupting ISR signaling via p-eIF2α haploinsufficiency or impaired *Atf4* expression reduces tumor burden and inhibits the formation of high-fitness clusters. (**a**) Representative ultrasound images of mouse lung tumors detected in the septum, located peripherally in contact with the pleura, were obtained 33 weeks after CRE-lentivirus intubation in eIF2α ^S/S^ (n = 7) and eIF2α ^S/A^ (n = 7) mice. Tumor is outlined with yellow intermittent lines. Data were analyzed using the Mann-Whitney U Test and are presented as mean ± SD. (**b**) The UMAP plot illustrates the distribution of tumor cells from eIF2α^S/S^ and eIF2α^S/A^ groups (n=15,588 cells), colored by annotated cell types. A density plot overlays the UMAP to visualize cell density across the latent space. (**c,d**) The scaled signature scores for the high stemness signature in panel C or the Hallmark MYC Target V1 signature in panel D were computed for all cell types in the eIF2α^S/S^, eIF2α^S/A^, and eIF2α^A/A^ models. The average score of single-cell transcriptomes within each group is displayed. (**e**) The distribution of Atf4(+) regulon activity scores, calculated using SCENIC, highlights the variation in Atf4-driven transcriptional activity across single cells. (**f**) Representative ultrasound images of mouse lung tumors in the septum, located peripherally in contact with the pleura, were obtained 33 weeks after CRE-lentivirus intubation (eIF2α^S/S^ Atf4**^+/+^**, n = 5 mice; eIF2α^S/S^ Atf4^+/-^, n = 5 mice). Tumor is outlined with yellow intermittent lines. Data were analyzed using the Mann-Whitney U Test and are presented as mean ± SD (**g**) The UMAP displays the distribution and density of tumor cells from eIF2α^S/S^ Atf4**^+/+^** and eIF2α^S/S^ Atf4^+/-^ groups (n = 19,943 cells), colored by annotated cell types. A density overlay highlights the concentration of cells across the UMAP space, while fractions illustrate the proportional representation of each cluster in the two genotypes.

We observed a shift in the placement of the high stemness cluster on the UMAP, reflecting a change in its gene expression profile in eIF2α^S/A^ compared to eIF2α^S/S^ tumors (Fig. 3b). Specifically, the high-stemness cluster in eIF2α^S/A^ tumors shows elevated expression of *Rbm3* (Extended Data Fig. 3e), a highly expressed marker gene of the Mito-Dysf cluster observed in eIF2α^A/A^ tumors (Fig. 2e and Supplementary Table 2). This increase is accompanied by a decreased expression of high-stemness cluster genes in eIF2α^S/A^ compared to eIF2α^S/S^ tumors (Fig. 3c and Supplementary Table 3). Collectively, these findings suggest that a partial reduction in p-eIF2α drives the high-stemness cluster toward adopting a Mito-Dysf fate, a feature typically associated with tumors exhibiting a complete loss of p-eIF2α.

Using the MYC transcriptional signature ^47^, which is an indicator of high stemness in eIF2α^S/S^ tumors (Fig. 1e and Supplementary Table 1), we found that the high stemness cluster of eIF2α^S/A^ tumors exhibits lower expression of MYC-dependent genes compared to eIF2α^S/S^ tumors (Fig. 3d). This difference in MYC upregulated gene expression suggests that partial ISR disruption in eIF2α^S/A^ tumors is enough to disrupt the acquisition of high stemness phenotype during KP lung tumor evolution.

We conducted single-cell regulatory network inference and clustering (SCENIC) ^48^ to identify transcriptional factors with a primary role in gene expression programs in each cluster. We found that ATF4 activity, as indicated by the expression of ATF4-dependent genes, is linked to the progression of lung transition cells to high stemness and pre-EMT clusters in eIF2α^S/S^ tumors (Figs. 2d and 3e). Conversely, ATF4 was not active in the transition of lung transition to Mito-Dysf cluster in eIF2α^A/A^ tumors (Figs. 2e and 3e). Since the Mito-Dysf cluster retains features of both AT2-like and AT1-like clusters, ATF4 activity in this cluster is sustained at its lowest level (Fig. 3e).

Considering the essential role of ATF4 downstream of p-eIF2α in the induction of ISR^49^, we investigated its role in lung tumor evolution in KP mice bearing a conditional *Atf4*^f/f^ allele. Heterozygous deletion of *Atf4^+/-^* specifically in KP lung tumors with intact p-eIF2α (eIF2α^S/S^) substantially decreased tumor growth (Fig. 3f), while homozygous deletion of *Atf4^-/-^* resulted in no detectable tumors by ultrasound imaging throughout the observation period.

Sc-RNA-seq analysis of the *Atf4^+/-^* lung tumors indicated the elimination of the high stemness, high plasticity and pre-EMT clusters (Fig. 3g). The remaining clusters in *Atf4^+/-^* tumors consisted of the AT2-like, lung transition, and gastric-like clusters, which were enriched compared to *Atf4^+/+^* tumors (Fig. 3g). These results suggest that the transition of KP tumors to undifferentiated late clusters strictly depends on the integrity of the eIF2α-ATF4 axis of ISR.

### The lung transition cluster is the epicenter of ISR-driven LUAD tumor evolution

We found that the AT2-like cells in the KP model evolve into the lung transition cluster (Fig. 2d), which serves as a central hub in the evolutionary process regulated by the ISR. This lung transition cluster resembles alveolar intermediate cells, specifically AT2-AT1 precursor cells, commonly found in embryonic lung development models and in models of lung injury and repair^29,50^. In these models, the ISR functions as a stress adaptation mechanism, allowing normal AT2 cells to transition into alveolar intermediate cells during lung regeneration or to support the development of the intermediate cell population^29,50^ However, persistent ISR activation can lead to an expansion of the AT2-AT1 precursor cell population, disrupting their normal differentiation process^29^.

When we examined normal mouse lung cell atlases^51–53^, focusing specifically on AT1, AT2, and AT2-AT1 intermediate cells (AT1/AT2), we observed that the AT2-like population within the KP tumors exhibits a gene expression profile closely resembling that of normal AT2 cells (Extended Data Fig. 4a). In the normal mouse lungs, AT2-AT1 intermediate cells form a bridge between the AT2, and lung transition populations seen in the tumors (Extended Data Fig. 4a). These data indicates that at early stages of KP tumor evolution, AT2-like and lung transition cluster display developmental characteristics of normal cells. Data analysis indicates that KRAS signaling, measured by the expression of ERK-dependent genes, is upregulated during lineage progression at a higher level in KP tumors compared to normal cells (Extended Data Fig. 4b).

We identified a specific subset of *KRAS-*mutated transitional cells, referred to as KRT8^+^ alveolar intermediate cells (KAC), which represent a state between normal AT2 and tumor cells ^22,54^. Our analysis revealed that KP tumor cells with similar expression profiles of human KAC exhibit elevated expression of ISR regulated pathways compared to alveolar transition cells (AICs) without *KRAS* mutations (Extended Data Fig. 4c). Furthermore, the KAC population in human LUAD closely mirrors the lung transition cluster observed in our KP model (Extended Data Fig. 4d). These findings underscore the relevance of our findings to human LUAD, as the human KAC population represents tumor-initiating cells ^22^.

The lung transition cluster follows two main trajectories: trans differentiation into gastric-like cells or dedifferentiation into high plasticity cells (Fig. 2d). Both pathways involve a reduction in the lung lineage factor *Nkx2-1*, a crucial regulator for maintaining lung cell identity and differentiation^55^. We observed a progressive decrease in *Nkx2-1* expression as eIF2α^S/S^ tumors dedifferentiated into late-stage clusters characterized by high plasticity and high stemness (Extended Data Fig. 4e and Supplementary Table 1). In contrast, eIF2α^S/A^ and *Atf4^+/-^* tumor cells exhibited only a modest reduction in *Nkx2-1* expression compared to the early stages (Extended Data Fig. 4e). Conversely, *Nkx2-1* expression remained high in eIF2α^A/A^ tumors, aligning with the transition of these tumors to the developmentally dead-end Mito-Dysf cluster (Extended Data Fig. 4e). Consistent with scRNA-seq data, IHC analysis showed a significant increase of NKX2-1 expression in eIF2α^A/A^ tumors compared to eIF2α^S/S^ tumors (Extended Data Fig. 4f). These results suggest that the ISR-mediated dedifferentiation of lung transition cluster in the KP model is at least partially driven by *Nkx2-1* downregulation.

### ISR-impaired tumor cells retain the dedifferentiation programs *ex vivo* mirroring primary tumors

Previous studies showed that *Itga2*, encoding integrin α2 (CD49B), a component of the integrin α2β1 collagen receptor, serves as a marker for human and mouse high plasticity cells^23,56^. We observed that *Itga2* expression is primarily evident in late-stage clusters, specifically in high plasticity, high stemness, and pre-EMT clusters in eIF2α^S/S^ tumors, and the Mito-Dysf cluster in eIF2α^A/A^ tumors (Fig. 4a). Based on scRNA-seq of eIF2α^S/S^ tumors, *Itga2*^high^ cells with characteristics of high plasticity, high stemness, and pre-EMT clusters accounted for approximately 30% of all tumor clusters (Fig. 4b).

**Fig. 4.**
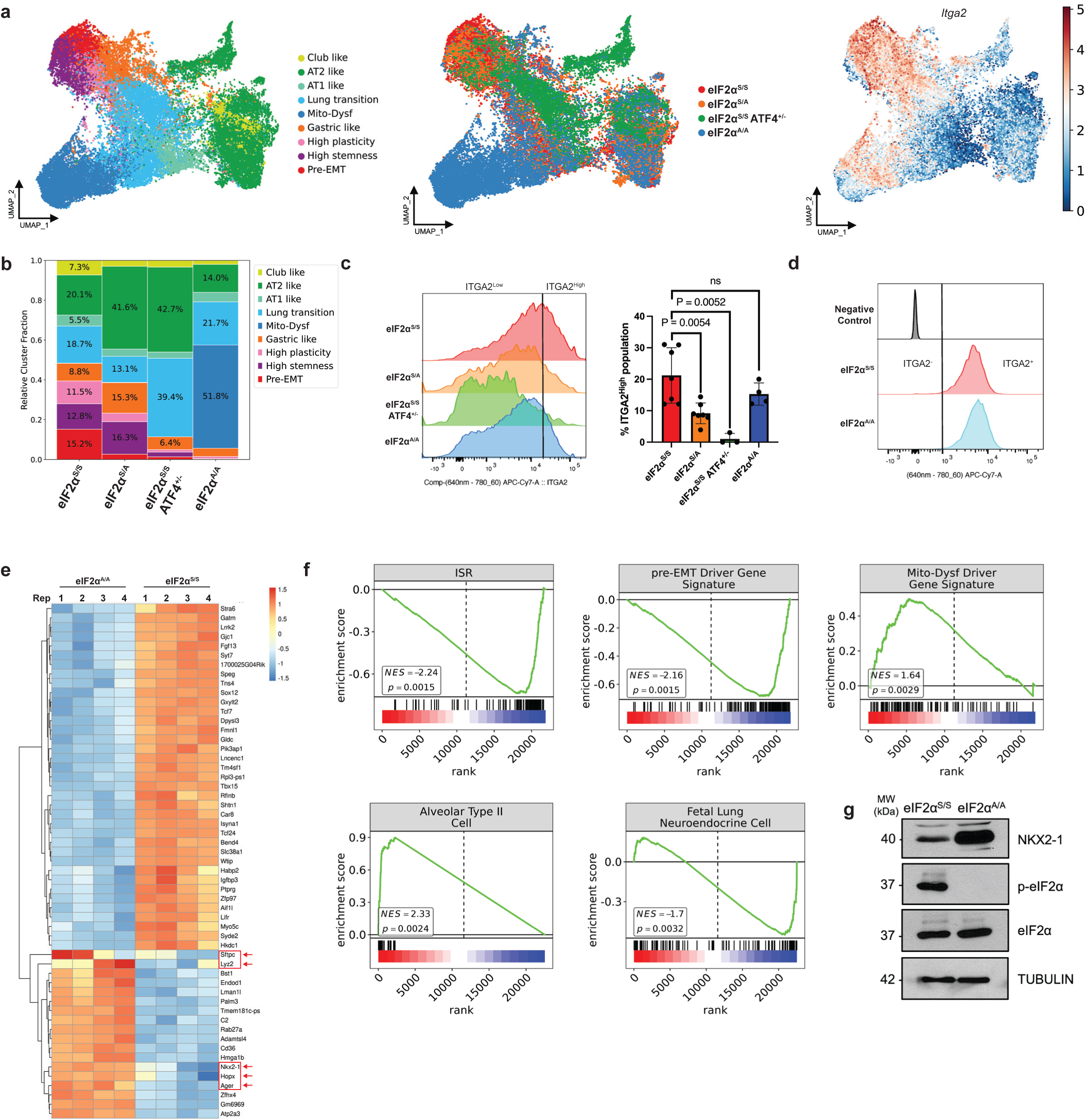
eIF2α^S/S^ and eIF2α^A/A^ KP cells cultured *ex vivo* preserve the transcriptional differences observed in primary tumors. (**a**) The UMAP plot represents the merged dataset (n = 44,780 cells) and is colored by cell types (left), genotype (middle), or Itga2 expression levels (right). (**b**) The bar plot illustrates cell type fractions grouped by genotype and colored according to the cell types specified in (**a**). (**c**) Histograms display the distribution of ITGA2 staining across the specified tumor types, analyzed by flow cytometry. The black dividing line indicates the threshold for high ITGA2 expression, defined as the top 30% of cells in the eIF2α^S/S^ group. The quantification of ITGA2^high^ cells, grouped by genotype, is presented on the right. Error bars represent the mean ± SD of biologically independent samples (eIF2α^S/S^, n = 7; eIF2α^S/A^, n = 7; eIF2α^A/A^, n = 3; eIF2α^S/S^ Atf4^+/-^, n = 4). Statistical significance was determined using the Mann–Whitney U-test, with p-values indicated in the graph. Results not reaching significance are labeled as “ns” (not significant) (**d**) Flow cytometry of ITGA2 staining of cultured eIF2α^S/S^ and eIF2α^A/A^ tumor cells. The fluorescence-minus-one (FMO) control was used as the negative control to determine background fluorescence. (**e**) Heatmap displays the top differentially expressed genes (DEGs) between eIF2α^S/S^ and eIF2α^A/A^ cultured cells. Arrows highlight alveolar marker genes on the heatmap, emphasizing transcriptional differences in alveolar cell identity between the two tumor genotypes. (**f**) GSEA was performed on pre-EMT, Mito-Dysf, ISR, fetal lung neuroendocrine, and AT2 cell transcriptional signatures to compare eIF2α^S/S^ and eIF2α^A/A^ tumor cells in culture. Negative enrichment scores indicate pathways enriched in eIF2α^S/S^ cells, while positive scores indicate enrichment in eIF2α^A/A^ cells. (**g**) Western blot analysis of the indicated proteins in eIF2α^S/S^ and eIF2α^A/A^ cultured cells.

Flow cytometry analysis confirmed that primary eIF2α^S/S^ KP tumors, freshly isolated from mouse lungs, displayed the highest proportion of ITGA2^high^ cells, followed by eIF2α^A/A^ tumors, while eIF2α^S/A^ and ATF4^+/-^ tumors had the lowest, aligning with the *Itga2* expression pattern across these groups (Fig. 4c). The impaired dedifferentiation in eIF2α^S/A^ and ATF4^+/-^ tumors, compared to eIF2α^S/S^ and eIF2α^A/A^ tumors that progress to late-stage developmental clusters characterized by high plasticity/stemness/pre-EMT and Mito-Dysf, respectively, likely explains the observed differences in ITGA2 levels (Fig. 4c). Therefore, ITGA2 levels can serve as a valuable marker for distinguishing tumors capable of advancing through developmental processes from those that cannot.

We transitioned from studying mouse tumors to cultured cells, developing a model to better understand the role of ISR in mitochondrial function in KP tumors. Flow cytometry analysis of cultured eIF2α^S/S^ and eIF2α^A/A^ cells indicated ITGA2^high^ expression in both tumor types (Fig. 4d), indicating that the cultured cells retained the late-stage characteristics of the primary tumors. This observation was further substantiated by RNA-seq analysis of the cultured tumor cells, showing upregulation of 1,702 genes and downregulation of 1,620 genes in eIF2α^S/S^ compared to eIF2α^A/A^ cells (fold change > 2, adjust P-value < 0.05). Differentiated alveolar cell markers such as *Nkx2-1*, *Hopx*, *Sftpc*, *Ager*, and *Lyz2* were more highly expressed in eIF2α^A/A^ cells compared to eIF2α^S/S^ cells (fold change > 2, adjust P-value < 0.05), indicating that cultured cells lacking p-eIF2α had the characteristics of KP tumor cells with an impaired evolutionary trajectory (Fig. 4e).

Our GSEA analysis indicates that eIF2α^S/S^ cells in culture predominantly exhibit a terminally dedifferentiated cell state with high stemness and pre-EMT characteristics, as inferred from the Cellrank fate-driving gene set (Fig. 4f and Supplementary Table 2). Additionally, pathway enrichment analysis shows that eIF2α^A/A^ cells have higher expression of the AT2 markers (Fig. 4f and Supplementary Table 2), whereas eIF2α^S/S^ cells display elevated expression of neuroendocrine cell markers (Fig. 4f and Supplementary Table 2). Furthermore, NKX2-1 expression is significantly lower in eIF2α^S/S^ compared to eIF2α^A/A^ cells in culture (Fig. 4g), aligning with sc-RNA seq data (Extended Data Fig. 4e) and IHC analysis of KP tumors (Extended Data Fig. 4f). These findings underscore that the phenotypic differences between eIF2α^S/S^ and eIF2α^A/A^ KP tumors are maintained even when these cells are cultured *ex vivo*.

### Tumor cells deficient in p-eIF2α display the Mito-Dysf phenotype *ex vivo*

Given that cultured eIF2α^S/S^ and eIF2α^A/A^ tumor cells preserve the evolutionary patterns observed in primary tumors, we investigated the role of the ISR in mitochondrial function within these cells. Transmission electron microscopy (TEM) analysis revealed a higher mitochondrial count in eIF2α^A/A^ compared to eIF2α^S/S^ cells (Fig. 5a), which aligns with the upregulation of genes encoding mitochondrial proteins in Mito Dysf cluster (Fig. 2g). However, despite the greater mitochondrial number, eIF2α^A/A^ cells displayed reduced oxygen consumption rates, including lower basal respiration, ATP-linked respiration, maximum respiration, and spare respiratory capacity (Fig. 5b), indicative of defective mitochondrial function. Furthermore, loss of p-eIF2α in eIF2α^A/A^ cells showed lower extracellular acidification rates (ECAR), reflecting a broader metabolic dysfunction (Fig. 5b).

**Fig. 5.**
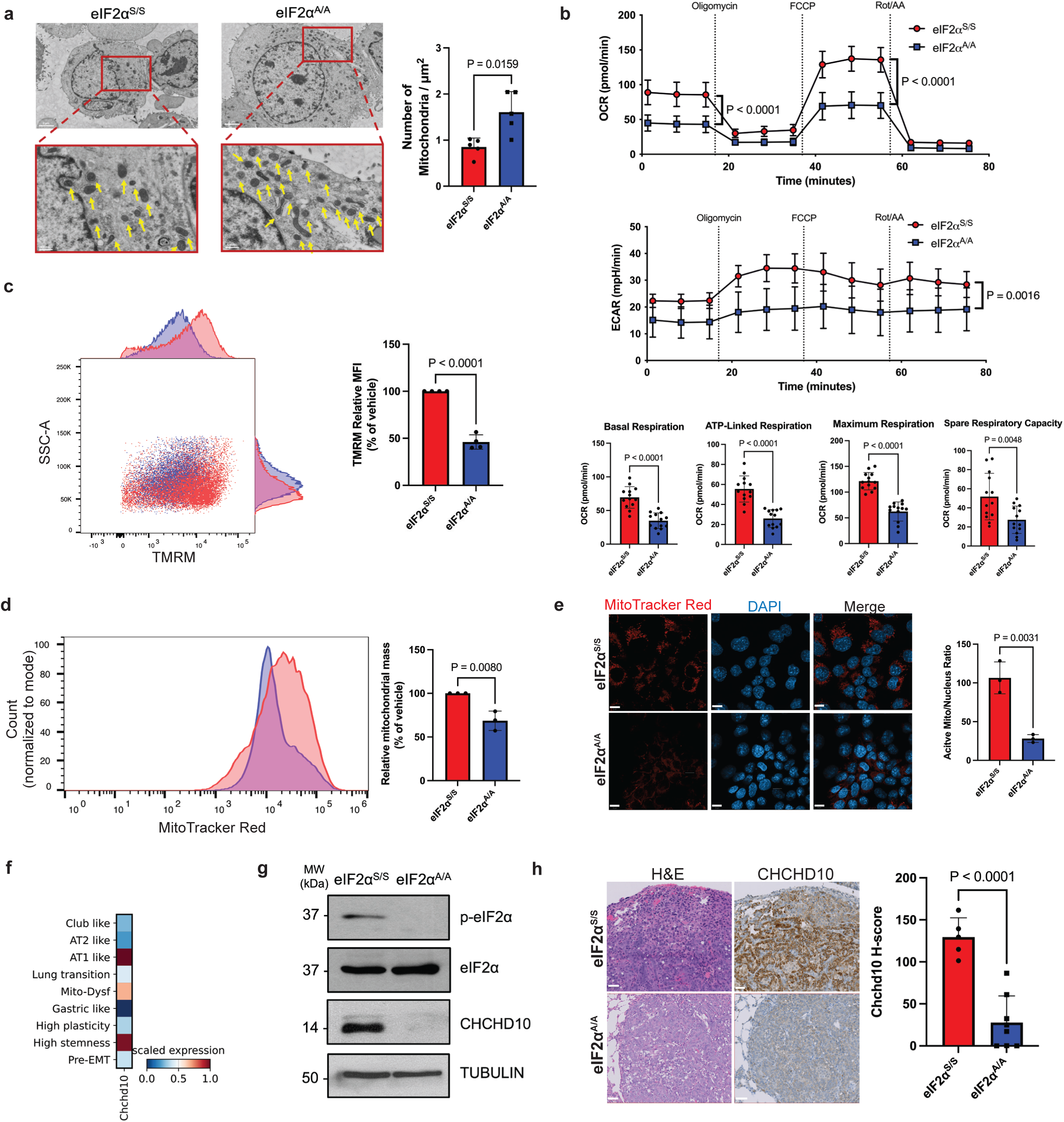
Tumors deficient in p-eIF2α exhibit a unique state of mitochondrial dysfunction, marked by disrupted mitochondrial processes. (**a**) Representative TEM images of mitochondria in cultured eIF2α^S/S^ and eIF2α^A/A^ tumor, with quantification based on the number of mitochondria per square micron. Scale bars represent 2 µm for full cell images and 500 nm for enlarged views. Yellow arrows indicate mitochondria. Quantification is based on n = 5 cells per tumor genotype. (**b**) Mitochondrial function was assessed in cultured eIF2α^S/S^ and eIF2α^A/A^ cells using the Seahorse assay, evaluating the oxygen consumption rate (OCR) and extracellular acidification rate (ECAR). Parameters measured include basal respiration, ATP-linked respiration, maximal respiration, and spare respiratory capacity. Data are presented as mean ± SD from n = 3 biological replicates, each performed in triplicates. (**c**) Mitochondrial membrane potential was measured in cultured eIF2α^S/S^ and eIF2α^A/A^ cells using TMRM fluorescence. Representative FACS plots display TMRM fluorescence levels, with quantification of mean fluorescence intensity expressed as a percentage relative to the vehicle control. Data are based on n = 4 biological replicates. (**d**) Active mitochondrial mass in cultured eIF2α^S/S^ and eIF2α^A/A^ cell lines was assessed by measuring the mean fluorescence intensity of MitoTracker using flow cytometry. Data are based on n = 3 biological replicates. (**e**) Confocal microscopy images of eIF2α^S/S^ and eIF2α^A/A^ cells in culture show active mitochondria stained with MitoTracker and nuclei counterstained with DAPI. The mitochondrial activity ratio was calculated by dividing the number of active mitochondria by the number of DAPI-stained nuclei. Scale bar = 10 µm. Data represent n = 3 biological replicates, each performed in triplicates (**f**) Heatmap shows *Chchd10* expression in different KP tumor clusters. (g) Western blot analysis of the indicated proteins. (h) IHC staining of CHCHD10 in eIF2α^S/S^ and eIF2α^A/A^ KP lung tumors; scalebars, 50 μm. (**a-e**) Data are presented as mean ± SD, with statistical analyses performed using a two-tailed unpaired t-test. P-values are displayed on the graph to indicate the significance of the results.

The mitochondrial dysfunction caused by the loss of p-eIF2α was further confirmed by a decrease in mitochondrial membrane potential in eIF2α^A/A^ compared to eIF2α^S/S^ cells (Fig. 5c). Additionally, eIF2α^A/A^ display a reduced mitochondrial activity, as evidenced by lower mitochondrial mass, confirmed through MitoTracker Red staining, and analyzed via flow cytometry (Fig. 5d) and confocal microscopy (Fig. 5e).

We observed elevated expression of the coiled-coil-helix-coiled-coil-helix domain containing 10 (*Chchd10*) gene in the high stemness cluster of eIF2α^S/S^ tumors compared to Mito-Dysf cluster of eIF2α^A/A^ tumors (Fig. 5f). *Chchd10* encodes a protein essential for maintaining inner mitochondrial membrane integrity and regulating mitochondrial respiration ^57^. Both in culture and in tumors, CHCHD10 expression was significantly impaired in eIF2α^A/A^ tumors compared to eIF2α^S/S^ tumors, as confirmed by immunoblotting and IHC (Fig. 5g,h).

These findings underscore the role of the ISR in safeguarding mitochondrial integrity and function in KP tumor cells.

### Inhibiting ISR pharmacologically halts tumor evolution and reduces growth

We further assessed the impact of the ISR inhibitor ISRIB, a small molecule that counteracts the translational effects of p-eIF2α ^3^, on LUAD progression in KP mice. After 35 weeks of tumor progression, mice with KP lung tumors and intact p-eIF2α (eIF2α^S/S^) were treated with either a vehicle control or ISRIB for a duration of 4 weeks. Treatment with ISRIB led to a significant reduction in tumor growth, as measured by ultrasound imaging (Fig. 6a).

**Fig. 6.**
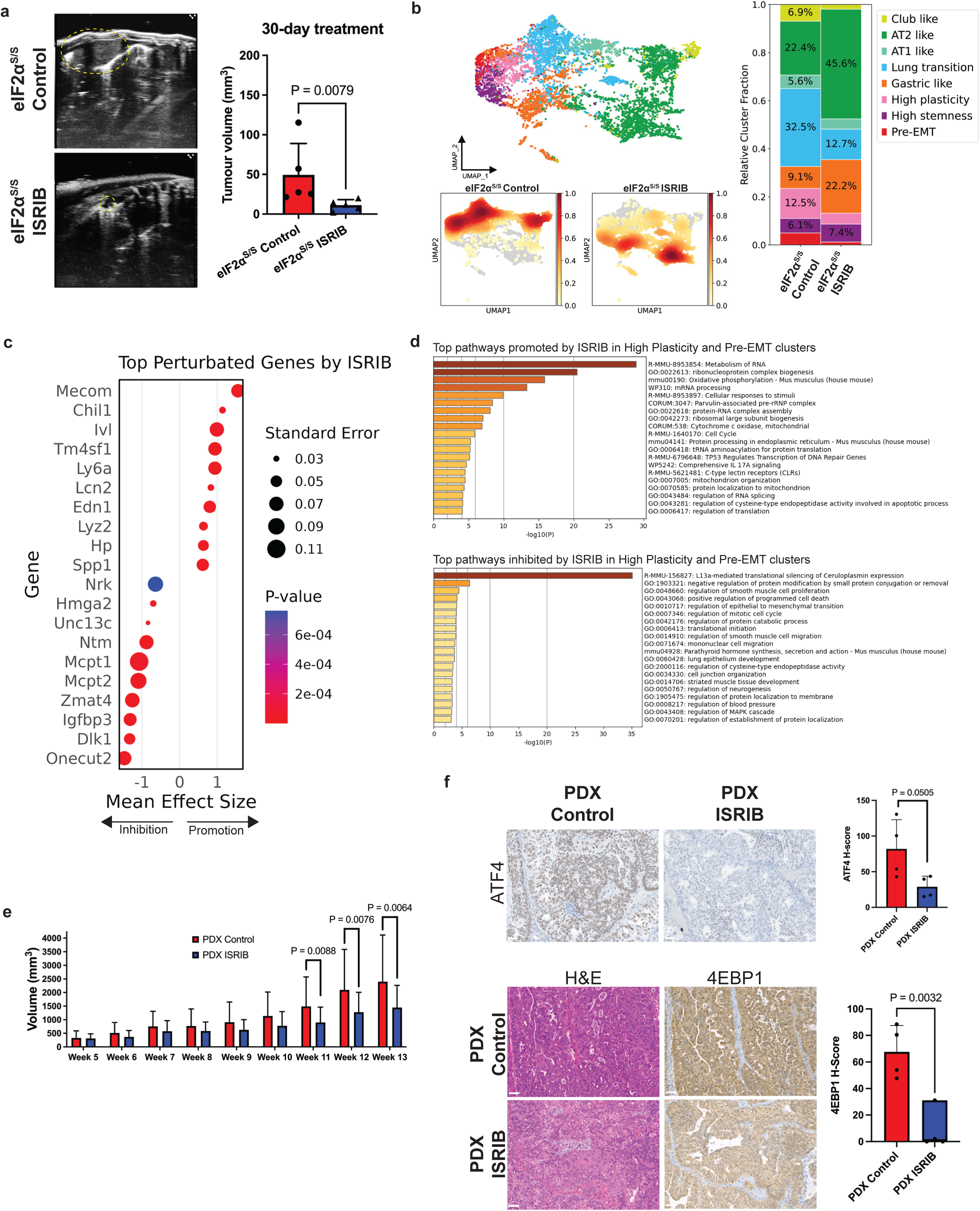
ISRIB treatment disrupts tumor evolution and primes tumors for the development of a mitochondrial dysfunction phenotype. (**a**) Representative ultrasound images of ISRIB-treated mouse lung tumors show detection in the septum, located peripherally in contact with the pleura, following 30 days of treatment administered 35 weeks after CRE-lentivirus intubation (eIF2α^S/S^ control treated n = 5 mice; eIF2α^S/S^ ISRIB-treated, n = 5 mice). Tumor is outlined with yellow intermittent lines. Data represent mean ± SD and were analyzed using the Mann-Whitney U Test. (**b**) The UMAP plot visualizes the distribution and density of tumor cells from eIF2α^S/S^ control and ISRIB treated groups (n = 8,580 cells), colored by annotated cell types. The density overlay highlights cell concentrations across the UMAP space, while cell type fractions illustrate the proportional representation of each annotated cell type in the two conditions. (**c**) The dot plots display the genes whose expression is most significantly affected by ISRIB treatment in eIF2α^S/S^ tumors. Dot size represents the proportion of cells expressing each gene, while color intensity indicates the average expression level. (**d**) Enrichment analysis was performed on ISRIB-affected genes, categorizing them into upregulated and downregulated groups. (**e**) ISRIB inhibits the subcutaneous growth of PDX LUAD tumors in immune-deficient NOG mice. Treatments began in week 5 following subcutaneous PDX transplantation. Mice were administered daily oral doses of either vehicle (n = 20 mice) or 10 mg/kg ISRIB (n = 20 mice). (**f**) H&E staining and IHC analysis were performed to assess nuclear ATF4 and cytoplasmic 4EBP1 expression in vehicle-control and ISRIB-treated PDX tumors after 4 weeks of treatment; scalebars, 50 μm. Quantification of ATF4 expression was conducted using samples from n = 3 mice per group, while 4EBP1 expression was quantified from n = 4 mice per group.

ScRNA-seq analysis revealed that ISRIB treatment suppressed the expansion of lung transitional cells and late-stage tumor cell populations, including high plasticity and pre-EMT clusters (Fig. 6b). Additionally, there was a marked increase in the AT2-like cell population within the residual tumor following ISRIB treatment (Fig. 6b). Flow cytometry analysis of KP eIF2α^S/S^ tumors confirmed that ISRIB treatment reduced ITGA2^high^ cells, shifting the distribution toward ITGA2^low^ cells (Extended Data Fig. 5a,b).

After ISRIB treatment, the AT2 marker *Lcn2* and mitochondrial marker *Atp5k* were significantly upregulated in all clusters, while the gastric regulator *Hnf4a* in gastric clusters showed a marked decrease (Extended Data Fig. 5c). Additionally, *Hmga2* expression was notably reduced in pre-EMT and high-plasticity clusters (Extended Data Fig. 5c). Overall tumor perturbation revealed the AT2 marker *Lcn2* as one of the most upregulated gene and pre-EMT marker *Onecut2* as the most downregulated gene post-treatment (Fig. 6c). Gene Ontology analysis highlighted enrichment of mitochondrial component genes linked to the Mito-Dysf fate in late-transition high-plasticity and pre-EMT clusters, accompanied by a reduction in mRNA silencing pathways following ISRIB treatment (Fig. 6d and Supplementary Table 4). However, the Mito-Dysf cluster was absent in scRNA-seq data, likely due to partial ISR inhibition by ISRIB alone. The level of *Onecut2, Hnf4a* and *Eif4ebp1* before and after ISRIB treatment indicates that the effectiveness of the treatment to target the late-stage tumor population, as these genes showed decreased expression following treatment (Fig. 6c, Extended Data Fig. 5d and Supplementary Table 4). The ISRIB phenotype was stronger than eIF2α^S/A^ tumors but weaker than in ATF4^+/-^ tumors (Extended Data Fig. 5e). In summary, ISRIB-treated tumors displayed phenotypic shifts and suppression of key evolutionary driver genes, indicating ISRIB’s ability to block the lineage plasticity driven by ISR and promote a Mito-Dysf-like phenotype over time.

We further investigated the impact of ISRIB on the growth of a human LUAD patient-derived xenograft (PDX) with a KRAS G12C mutation in immunodeficient NOG mice (Fig. 6e). Daily oral administration of ISRIB for 13 weeks resulted in approximately 30% inhibition of PDX tumor growth compared to vehicle-treated tumors (Fig. 6e). IHC analysis of tumor sections revealed reductions in 4EBP1 and nuclear ATF4, which both are markers of late-stage, dedifferentiated tumors (Fig. 6f).

Overall, the data demonstrates that ISRIB suppresses tumor growth and dedifferentiation markers in human LUAD models, highlighting its potential as a therapeutic strategy for KRAS-mutant lung cancers.

### ISR is a marker of evolution and a potential therapeutic target of human LUAD tumors

Analysis of gene expression profiles from 539 LUAD tumor samples in The Cancer Genome Atlas (TCGA) and 706 non-mucinous LUAD tumor samples from surgically treated patients revealed that ISR-dependent gene expression, along with high plasticity, pre-EMT, and high-stemness programs, is associated with poorer survival outcomes (Fig. 7a,b, Extended Data Fig. 6 and Supplementary Table 5). In contrast, tumors characterized by high expression of AT2-like, AT1-like, and lung transition profiles were linked to longer overall survival (Fig. 7a,b, Extended Data Fig. 6 and Supplementary Table 5). Notably, this trend also holds true for LUAD patients harboring *KRAS* mutations (Fig. 7c and Supplementary Table 5).

**Fig. 7.**
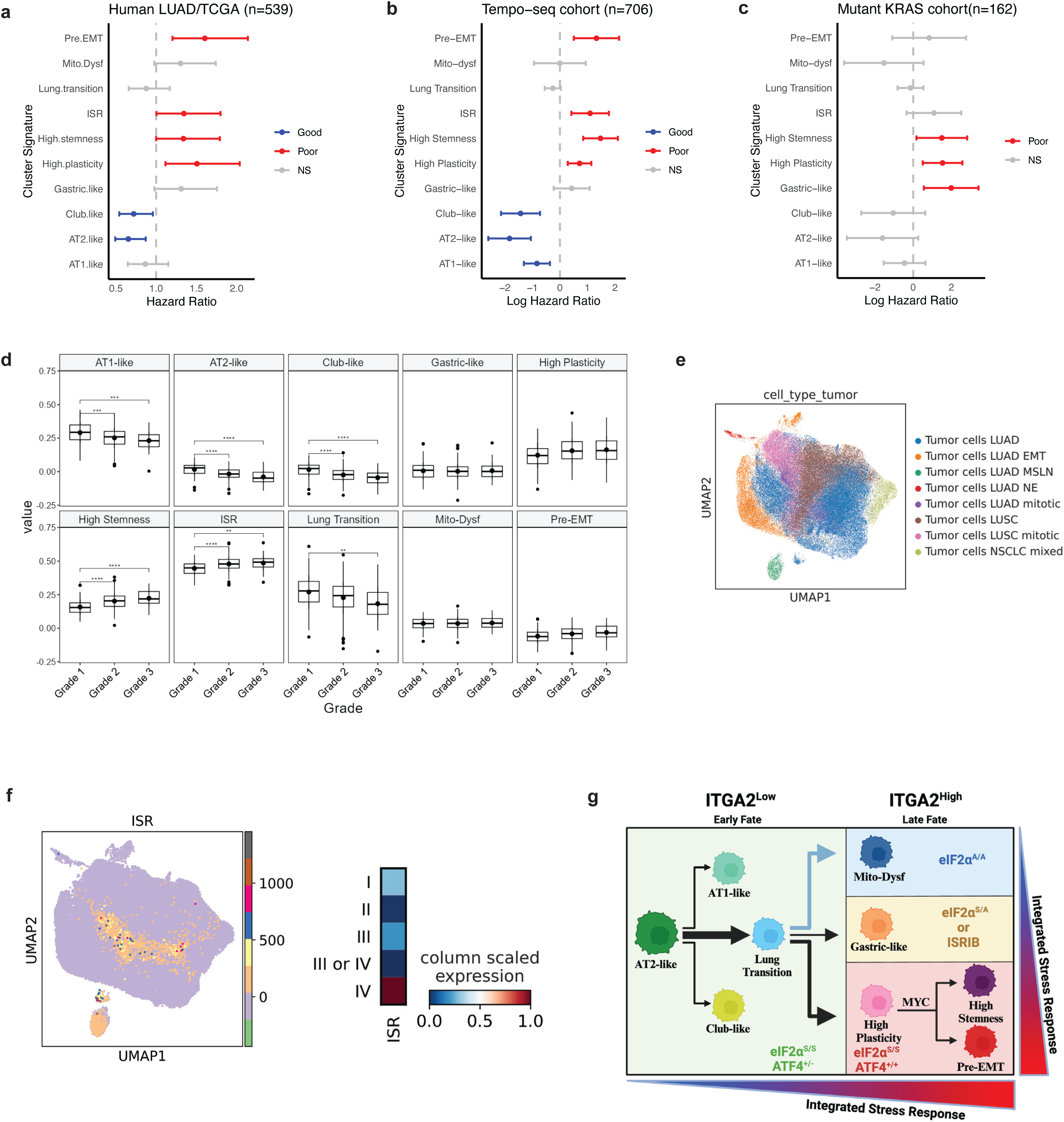
ISR signaling and ISR-driven dedifferentiated tumor cells are present in human LUAD and correlate with poor patient survival. (**a-c**) The hazard ratios (HRs) for cluster markers in LUAD patient samples were calculated using Cox proportional hazards models, with each cluster marker activity term treated as a continuous variable. The x-axis represents HRs, including their mean values and 95% confidence intervals. Panel A: HRs for LUAD patient samples in TCGA database (*n=*539). Panel B: HRs for the entire LUAD patient cohort (*n* = 706). Panel C: HRs for the LUAD patient cohort with KRAS mutations (*n* = 162). (**d**) Box plots showing the expression levels of LUAD cluster signatures across different tumor grades. Statistical significance is indicated on the Fig. (*p < 0.05, **p < 0.01, ***p < 0.001). (**e,f**) The UMAP plots provide a visual representation of human lung cancer patient tumor samples: Panel E: Tumor cells are grouped and colored based on lung cancer cell classifications, highlighting the diversity of cell types within the TME. Panel F: Tumor cells are colored to indicate the intensity of the ISR transcriptional signature, showcasing the distribution and variation of ISR activity across different cells. The heatmap depicting scaled ISR expression scores across various Union for International Cancer Control (UICC) stages. The analysis includes 83,439 cells, illustrating the dynamic changes in ISR activity throughout LUAD progression. (**g**) Model of regulation of LUAD tumor evolution by ISR: LUAD tumor evolution begins with AT2-like cells differentiating into various clusters with low ISR activity. Gradual ISR activation via p-eIF2/ATF4 drives dedifferentiation into high-plasticity, high-stemness, and pre-EMT clusters. Partial p-eIF2 loss (eIF2α^S/A^) or ISRIB treatment restricts dedifferentiation to a gastric-like cluster with mitochondrial dysregulation, while complete loss (eIF2α^A/A^) results in a mitochondrial dysfunctional cluster (Mito-Dysf). The p-eIF2/ATF4 pathway, together with MYC, drives late-stage high-plasticity and high-stemness clusters, with ISR programs differing between stemness and pre-EMT clusters. The schematic illustrating this model was created using BioRender (https://www.biorender.com)

Our analysis further revealed varying degrees of ISR dependence across different cell populations within LUAD patient samples. Notably, cell populations associated with favorable prognosis, characterized by signatures such as AT1-like, AT2-like, club-like, and lung transition, showed a marked decline in expression as tumor grade increased (Fig. 7d). Conversely, cell populations with poor prognosis signatures, including those with high stemness and elevated ISR activity, demonstrated a significant increase in expression with advancing tumor grade (Fig. 7d).

We further analyzed datasets compiled into a comprehensive scRNA-seq atlas of lung cancer, encompassing 232 NSCLC patients and 86 non-cancer controls (Fig. 7e) ^58^. Using the computational tool Scanpy for scRNA-seq analysis, we identified a significant elevation of ISR activity in stage 4 human tumors (Fig. 7f). The analysis involved clustering cells based on their transcriptional profiles, which revealed distinct ISR-associated gene expression signatures prominently upregulated in advanced-stage tumors. The findings highlight the role of the ISR in driving cellular state changes and tumor progression in human LUAD, mirroring findings in the mouse KP model.

## DISCUSSION

### ISR’s role in LUAD progression and tumor plasticity

The study examines the role of the ISR in LUAD progression, particularly the transition from tumor-initiating cells to advanced, stem-like cells. Single-cell transcriptome analysis of a genetically engineered mouse model revealed that elevated ISR activity correlates with the emergence of high-plasticity, high-stemness cell populations, providing insights into how LUAD cells adapt under stress (Fig. 7g).

Our findings demonstrate that the ISR promotes tumor dedifferentiation into highly plastic cells, which are pivotal in creating high-stemness and pre-EMT trajectories that contribute to the heterogeneity of advanced-stage cancers^23^. In our model, the high plasticity cells are crucial for the formation of high stemness and pre-EMT cell fate trajectories stimulated by ISR (Fig. 2d,h). Both the high stemness and pre-EMT clusters display elevated levels of stemness (Fig. 2h), though the pre-EMT cluster shows a more pronounced tendency toward neuroendocrine cell characteristics (Fig. 1d, Extended Data Fig. 1a and Supplementary Table 1). Cells with neuroendocrine characteristics are known to play a pivotal role during the histological transformation of EGFR-driven LUAD into small cell lung cancer (SCLC)^25^.

### The interplay between MYC and ATF4 determines ISR-mediated tumor cell dedifferentiation

In EGFR-driven LUAD models, tolerance to MYC activity—a driver of pulmonary neuroendocrine lineage^59^—is essential for histological transformation (HT) into SCLC^25^. The integration of the single-cell transcriptomics data from the EGFR-driven LUAD model and our KP model indicates that precursor cells to SCLC correspond to the high-stemness cluster in our model (Extended Data Fig. 7a) and exhibit elevated ISR expression (Extended Data Fig. 7b). These findings suggest that ISR aids cancer cells in dedifferentiation and managing MYC-induced stress, reinforcing its oncogenic role in LUAD and potentially its involvement in SCLC transformation. Further research is needed to elucidate this mechanism.

Our study shows that the ISR and MYC pathways are co-upregulated in high plasticity, high stemness, and pre-EMT clusters during late evolutionary stages of tumor progression (Fig. 1e and Supplementary Table 1). As lung cell identity diminishes, cells increasingly depend on MYC, rather than mutant KRAS, to maintain stem cell properties (Fig. 1e) ^25,60^. MYC upregulation in the high plasticity population may also enhance ISR activity by activating stress-related eIF2α kinases like GCN2 and PERK^49^. This link may explain the elevated ISR signaling in these clusters.

We observed that the poorly differentiated clusters exhibit distinct MYC-dependent oncogenic pathways. The high stemness cluster exhibits the highest MYC-dependent gene expression and upregulated mTOR signaling (Fig. 1e and Supplementary Table 1), which can enhance ATF4 expression and fine-tune MYC-driven translational effects^49,61,62^.

In contrast, the high plasticity and pre-EMT clusters rely more on p-eIF2α for mRNA translation regulation due to lower MYC activity and reduced mTOR signaling (Fig. 1e and Supplementary Table 1), which can lead to increased p-eIF2α levels^63^. These clusters are particularly sensitive to p-eIF2α deficiency, as seen in tumors with haploinsufficient p-eIF2α, where high plasticity and pre-EMT populations are significantly reduced (Fig. 3b). Future studies will explore whether ATF4 expression during ISR mirrors hormonal *Atf4* gene regulation^64,65^.

ATF4 is essential for MYC-driven tumor progression by regulating genes involved in amino acid influx and protein synthesis^49^. Loss of ATF4 (*Atf4*^+/-^) halts the progression of cells in the lung transition cluster, preventing their development into high stemness, pre-EMT, or high plasticity clusters, and abolishes MYC-dependent gene expression (Fig. 3g). The ATF4-MYC collaboration prevents protein toxicity by upregulating 4E-BP1^49,66^, ensuring controlled mRNA translation and supporting LUAD progression.

The upregulation of 4E-BP1, which inhibits cap-dependent translation, supports cell survival under nutrient and oxidative stress^66^, prevents TP53-dependent senescence while facilitating oncogene-driven transformation of primary fibroblasts^67^, maintains mitochondrial homeostasis to avert mesenchymal stem cell senescence^68^, and acts as a metabolic switch during glucose starvation^69^. In LUAD, our data suggests that 4E-BP1 could potentially drive tumor evolution toward high stemness and pre-EMT clusters, highlighting the importance of the MYC-ATF4 axis in tumor progression (Fig. 2e,f). High 4E-BP1 expression, observed in nearly all 33 cancer types in TCGA, including LUAD, is associated with poor prognosis, underscoring its critical role in cancer biology^70^.

### Mitochondrial dysfunction as a barrier to tumor evolution in KP LUAD model

Mitochondrial dysfunction is a defining feature of KP tumor clusters, particularly in eIF2α^A/A^ and *Atf4^+/-^* KP tumors, where the absence of both p-eIF2α and ATF4 impairs tumor evolution. In eIF2α^A/A^ tumors, most cells remain in the Mito-Dysf state, characterized by incresed *Nkx2-1* expression levels similar to the lung transition cluster (Fig. 1d), indicating an inability to complete dedifferentiation into high-plasticity cells essential for advanced tumor progression and therapy resistance^23^.

The Mito-Dysf cluster exhibits upregulation of genes linked to mitochondrial ribosomal biogenesis and oxidative phosphorylation (Figs. 1e and 2g and Supplementary Table 1), critical for metabolism, growth, and proliferation^71^. Analysis of KP cells *ex vivo* revealed disrupted mitochondrial membrane potential and respiration due to p-eIF2α loss (Fig. 5b-e), with an increased mitochondrial population potentially compensating for metabolic stress caused by dysfunction (Fig. 5a).

The loss of p-eIF2α may impair mitochondrial function by downregulating CHCHD10 expression (Fig. 5f-h), a critical factor in mitochondrial respiration and cellular adaptation to mitochondrial stress ^72^. Given that mutations in *Chchd10* are linked to neurological disorders ^57^, our data suggest a broader role for CHCHD10 in human diseases, including LUAD tumor progression.

### Targeting the ISR to impair LUAD progression and tumor growth

Cell plasticity enables tumor cells to adapt to stress and evade treatments, contributing to therapy resistance in cancers like LUAD ^73^. Chemotherapy in KRAS-mutant LUAD promotes high-plasticity, therapy-resistant neuroendocrine phenotypes ^74–77^. The dedifferentiation of LUAD into resistant clusters depends on the ISR pathway, which correlates with poor survival and advanced cancer stages (Fig. 7). Elevated p-eIF2α levels are markers of poor prognosis for LUAD patients^11^, and ISR-driven dedifferentiation is observed in both KRAS- and EGFR-driven LUAD tumors (Extended Data Fig. 7).

Targeting the ISR is a promising strategy to enhance cancer treatment sensitivity ^11,17,78^. ISRIB, a small molecule that counteracts p-eIF2α’s translational effects ^3^, significantly reduces tumor growth and extends the lifespan of mice in KRAS-driven LUAD models without detectable side effects, even after prolonged use^11^. ISRIB exerts its anti-tumor effects by reducing late-stage clusters, which are linked to therapy resistance^23,73^, while priming them toward a Mito-Dysf phenotype (Fig. 6c,d). Its ability to cross the blood-brain barrier and ongoing clinical trials for neurodegenerative diseases suggest its potential for repurposing in LUAD and other ISR-dependent cancers ^3^.

In the PDX LUAD model, ISRIB effectively impaired tumor growth (Fig. 6e); however, its efficacy was lower compared to its suppression of tumor development in the KP mouse model with an intact immune system (Fig. 6a). This difference may be due to the absence of a functional anti-tumor immune response in the PDX model, as mitochondrial stress responses and the ISR are linked to cancer immunity regulation ^79^.

Mutant KRAS inhibitors disrupt LUAD cell dedifferentiation in mice, leading to AT1-like cells capable of differentiating into tumorigenic clusters ^80^ and residual subpopulations expressing *Onecut2* and *Hnf4a* genes ^81^. ISRIB suppresses key pre-EMT markers (Fig. 6c), highlighting its potential to target KRAS-resistant subpopulations and improve therapeutic outcomes. Our findings propose ISRIB as a promising therapy to block dedifferentiation, reduce heterogeneity, and improve treatment outcomes in LUAD.

## METHODS

### Transgenic mouse model and treatment

Lung tumorigenesis was induced in KRAS^+/LSL-G12D^;fTg/0;eIF2α^S/S^ and KRAS^+/LSL-G12D^;fTg/0;eIF2α^A/A^ mice on C57BL/6 background through intratracheal intubation with lentiviruses expressing CRE and TP53 shRNA^11^. Lung tumor development was monitored using ultrasound imaging on the VisualSonics VEVO 3100 high-frequency ultrasound system^82^. To impair ATF4 in mouse LUAD tumors, KRAS^+/LSL-G12D^;fTg/0;eIF2αS/S mice were crossed with ATF4^f/f^ mice on a C57BL/6 background^49^. ISRIB treatment was administered daily by oral gavage, using a solution of 0.5% Hydroxypropylmethylcellulose (HPMC) and 0.1% Tween 80, pH 4.0, at a dose of 10 mg/kg^11^.

### Patient-derived xenograft (PDX) assay

The patient-derived xenograft (PDX) tumor model of LUAD carrying *KRAS G12C* was obtained from Jacksons labs (Tumor model: TM00186 LG0418F). The PDX model was established from a female 68 years-old Caucasian patient who was not subjected to chemotherapy, immunotherapy, hormone, or radiation therapy within 5 years prior to sample collection (treatment naïve). The PDX was amplified by subcutaneous transplantations in NOD.Cg-*Prkdc^scid^ Il2rg^tm1Wjl^*/SzJ (NSG) donor mouse from Jackson Laboratories according to provider’s specifications. Tumor growth in mice was measured with digital calipers two times per week, and the volume calculated by the formula: tumor volume [mm^3^] = [(length [mm]) × (width [mm])^2^]/2. The treatment with ISRIB was administered daily to mice by oral gavage, following the protocol used for the autochthonous model.

### Guidelines of ethical conduct in mouse work

The animal studies were performed in accordance with the Institutional Animal Care and Use Committee (IACUC) of McGill University and procedures were approved by the Animal Welfare Committee of McGill University (protocol #5754).

### Protein extraction and immunoblotting

Cells were washed twice with ice-cold PBS and proteins were extracted in ice-cold Radioimmunoprecipitation Assay (RIPA) buffer containing 10 mM Tris-HCl, pH 7.5, 50 mM KCl, 2 mM MgCl2, 1% Triton X-100, 3 μg/ml aprotinin, 1 μg/ml pepstatin, 1 μg/ml leupeptin, 1 mM dithiothreitol, 0.1 mM Na3VO4, and 1 mM phenylmethylsulfonyl fluoride. Cell lysates were kept on ice for 15 min, centrifuged at 13,200 x g for 15 minutes at 4 °C, and the supernatants were stored at −80 °C. Protein concentrations were measured using Bradford assay (Bio-Rad). The expression of different proteins was determined by loading 50 µg of the protein extracts from the same set of samples on 10 % sodium dodecyl sulfate (SDS)-polyacrylamide gels. After electrophoresis and protein transfer to Immobilon-P membrane (Millipore), blots were probed for phosphorylated eIF2α (Abcam, cat#Ab32157, 1:1000), total eIF2α (Cell Signaling Technologies, cat# L57A5, 1:1000), CHCHD10 (Proteintech, cat#25671-1-AP, 1:1000), NKX2-1 (Abcam, cat# EP1584Y, 1:2000), TUBULIN (Sigma-Aldrich, cat#T5168, 1:4000). Membranes were then probed with the corresponding secondary antibody: mouse IgG-horseradish peroxidase (HRP)-conjugated secondary antibody (KPL, cat#474-1806, 1:2000) for TUBULIN and rabbit IgG-HRP-conjugated secondary antibody (Cell Signaling Technologies, cat#7074, 1:1000) for the other primary antibodies. The protein expression was visualized by enhanced chemiluminescence (ECL, Thermo Fisher Scientific, cat#32106) according to the manufacturer’s specifications.

### Transmission electron microscopy (TEM)

Cells were plated 48 hours before the experiment to reach 80-90% confluency on the day of collection. Cells were detached using PBS-EDA and centrifuged at 300 g for 10 minutes. After discarding the supernatant, electron microscopy samples were fixed overnight in 2.5% glutaraldehyde in 0.2M cacodylate buffer (pH 7.2) at 4°C, followed by post-fixation in a 1:1 mixture of 2% osmium tetroxide and 2% potassium ferrocyanide in distilled water for 1 hour at 4°C with agitation. After post-fixation, samples were washed three times in distilled water for 5 minutes each at room temperature with agitation, then counter-stained with 1% uranyl acetate in distilled water for 30 minutes, protected from light. Samples were subsequently washed once in distilled water for 5 minutes at room temperature with agitation. Dehydration was carried out through a graded series of acetone or ethanol concentrations (25%, 50%, 75%, 95%, 100%, and 100%) for 30 minutes each at room temperature with agitation. For pre-infiltration, samples were incubated in graded ethanol/acetone to SPURR resin mixtures (3:1, 1:1, and 1:3) for 2 hours each at room temperature with agitation. This was followed by infiltration with pure SPURR resin for two intervals, first for 2 hours and then for an additional 3 hours, uncovered, at room temperature with agitation. Finally, samples were transferred to BEEM capsules and cured overnight at room temperature, then embedded at 60°C for 48 hours. After cooling briefly, blocks were demolded, ultrathin sections were cut with an ultramicrotome, collected Formvar Carbon 200 mesh copper grids (Sigma-Aldrich), stained with uranyl acetate and lead citrate, and imaged using a Hitachi H-7100 transmission electron microscope with AMT Image Capture Engine (version 600.147) at the INRS-CAFSB platform.

### Mitochondrial function assays

Mitochondrial membrane potential was assessed using TMRM staining in flow cytometry. Cells were incubated with TMRM at a final concentration of 20 nM for 30 minutes. After staining, cells were washed with PBS and analyzed by flow cytometry. Mean fluorescence intensity (MFI) was recorded, and mitochondrial membrane potential was quantified as a percentage relative to the vehicle control.

Mitochondrial content was assessed using Mitotracker Red staining for both flow cytometry and confocal microscopy. Cells were incubated with Mitotracker Red at a final concentration of 10 nM for 45 minutes. For flow cytometry, stained cells were analyzed immediately, with MFI used to quantify mitochondrial content. For confocal microscopy, cells were fixed post-staining and imaged using a Zeiss LSM 800 microscope to visualize mitochondrial distribution and morphology.

Mitochondrial function was assessed using a Seahorse XFe96 Analyzer (Agilent Technologies). Tumor cells were seeded in Seahorse XFe96 cell culture plates at a density of 4,000 cells per well 48 hours before the experiment in XF media (non-buffered RPMI containing 2 mM L-glutamine, 25 mM glucose, and 1 mM sodium pyruvate). Baseline OCR and ECAR were recorded with three basal measurements, followed by injections of mitochondrial inhibitors at the following concentrations: oligomycin (1 µM, Sigma-Aldrich), fluorocarbonyl cyanide phenylhydrazone (FCCP, 2 µM, Sigma-Aldrich), and a combination of antimycin A (0.5 µM, Sigma-Aldrich) and rotenone (0.5 µM, Sigma-Aldrich). Three OCR and ECAR measurements were taken after each injection. Basal respiration was calculated as the last basal OCR measurement minus the non-mitochondrial respiration rate after antimycin A/rotenone injection. ATP-linked respiration was determined by the reduction in OCR after oligomycin addition, while spare respiratory capacity was quantified as the absolute increase in OCR following FCCP injection relative to basal respiration. All measurements were normalized to cellular protein content, determined by sulforhodamine B (SRB) assay. Oxygen consumption and acidification curves were generated and analyzed using Agilent Seahorse Analytics software and GraphPad Prism 10.3.1.

### Immunohistochemistry (IHC)

Tissue samples were cut at 4-µm, placed on TOMO slides (VWR) and dried overnight at 37°C, before IHC processing. The slides were then loaded onto the Discovery XT Autostainer (Ventana Medical System). All solutions used for automated immunohistochemistry were from Ventana Medical System (Roche Tissue Diagnostics) unless otherwise specified. Slides underwent de-paraffinization, heat-induced epitope retrieval (CC1 prediluted solution, Roche Tissue Diagnostics, 06414575001). Immunostaining for ONECUT2, EIF4EBP1, CHCHD10, NKX2-1 was performed using a heat protocol. Briefly, rabbit polyclonal anti-ONECUT2 (Proteintech, cat# 21916-1-AP), anti-EIF4EBP1 (Cell Signaling Technology, cat#9452), anti-CHCHD10 (Proteintech, cat#25671-1-AP), and anti-NKX2-1 (Abcam, cat# EP1584Y) diluted at in the antibody diluent (Roche Tissue Diagnostics, 06440002001) were manually applied for 32min at 37°C then followed by the appropriate detection kit (OmniMap anti-Rabbit-HRP, Roche Tissue Diagnostics, 05269679001, for 8min, followed by ChromoMap-DAB, Roche Tissue Diagnostics, 05266645001). A negative control was performed by the omission of the primary antibody. Slides were counterstained with Hematoxylin (Roche Tissue Diagnostics, 05266726001) for 12 minutes, blued with Bluing Reagent (Roche Tissue Diagnostics, 05266769001) for 4 minutes, removed from the autostainer, washed in warm soapy water, dehydrated through graded alcohols, cleared in xylene, and mounted with Eukitt Mounting Medium (Electron Microscopy Sciences, 15320). Sections were analyzed by conventional light microscopy or scanned using the Aperio AT Turbo Scanner (Leica Biosystems). Mouse tissues were fixed in 10% buffered formalin phosphate, paraffin embedded, and sectioned. Paraffin was removed from the sections after treatment with xylene, rehydrated in graded alcohol, and used for H&E staining and immunostaining. Antigen retrieval was performed in sodium citrate buffer. Primary antibodies were incubated at 4 °C overnight and secondary antibodies were incubated at room temperature for 90 min. Sections were counterstained with 20% Harris modified hematoxylin (Thermo Fisher Scientific), mounted in Permount solution (Thermo Fisher Scientific), and scanned using a ZEISS Axioscan6. Quantification of stained sections was performed using QuPath 0.5.0.

### Isolation of mouse LUAD cells

Mice with LUAD tumors were euthanized 35 weeks post-tumor induction. Tumor samples were collected by isolating whole lungs, which were then dissociated using a solution of Dispase II (Gibco, cat#17105-041; 0.6 U/ml final concentration), Collagenase Type IV (Thermo Fisher Scientific, cat#17104019; 0.083 U/ml final concentration), and DNase I (Sigma-Aldrich, cat#69182-3; 10 U/ml final concentration) in RPMI 1640 (Wisent, cat# 350-700 CL. This mixture was incubated with the lungs at 37°C for 60 minutes. Dissociated cells were filtered through a 100 μm strainer and centrifuged at 300 g for 5 minutes at 4°C. After aspirating the supernatant, red blood cell lysis was performed using ACK Lysing Buffer (Thermo Fisher Scientific, cat#A1049201). The LUAD cells were then washed with media, pelleted at 300 g for 5 minutes at 4°C, and resuspended in Fluorescence-Activated Cell Sorting (FACS) buffer media (200 mM EDTA with 250 μl heat-inactivated FBS in PBS). Cells in suspension were incubated with appropriate stains for 20 minutes at 4°C, washed twice with ice-cold PBS, and resuspended in FACS buffer. Cells were then strained into FACS tubes using a 70 µm cell strainer before analysis on a BD FACSAria™ Fusion Flow Cytometer. FACS data were collected with FACSDiva software and analyzed using FlowJo software.

### Cell lines

Primary KRAS G12D eIF2α^S/S^ and eIF2α^A/A^ lung tumor cells were isolated from mice as previously described ^11^ and maintained in RPMI 1640, 10% FBS, antibiotics (100 units penicillin/streptomycin), 0.075% Sodium Bicarbonate NaHCO_3_ (Life Technologies), 1X essential amino acids (Life Technologies), 1X non-essential amino acids (Life Technologies). Due to the slow growth of KP eIF2α ^A/A^ and ATF4^+/-^ tumor cells, the primary cells were cultured until they reached adequate numbers for the preparation of frozen stocks.

### cDNA Library Preparation

Single-cell steps were performed using Chromium Next GEM Single Cell 3’ Kit v3.1 (10X Genomics). Briefly, sorted cells were manually counted and loaded on the Chromium instrument (10X Genomics). The target cell recovery was 8000 cells for tumor samples. Reverse transcription and library steps were performed; cDNA and library profiles were validated using a 2100 Bioanalyzer (Agilent). Final libraries were sequenced PE100 on a NovaSeq (Illumina).

### Bulk RNA-seq Analysis

Total RNA of KRAS G12D eIF2α^S/S^ and eIF2α^A/A^ cells (four replicates each) was isolated with Trizol (Thermo Fisher Scientific) and RNA-Seq libraries were prepared following the TruSeq Stranded Total RNA protocol (Illumina) according to the manufacturer’s instructions and 50 base single-end reads were obtained using a HiSeq2500 system in Rapid Mode (Illumina). FASTQ files were processed using the standard DESeq2 pipeline (version 1.24.0). Reads were aligned to the GRCm38 reference mouse genome. Count matrices were generated using DESeq2. Data were filtered to only include genes with 10 or more counts in 2 or more samples. The significance of differential expression between vehicle and treatment groups was quantified by one-way ANOVA. Genes with absolute Fold Change (FC) > 2 and False Discovery rates (FDRs) < 0.05 were considered differentially expressed. Gene set enrichment analysis (GSEA v4.0.3, Broad Institute) was performed on all genes ranked according to fold change, using the Gene Ontology geneset v5.2 (MSigDB)^47,83,84^. The number of permutations was 1000 and only sets containing between 15 and 500 genes were retained.

### Single-cell (sc) RNA sequencing

FASTQ files of scRNA-seq data generated on the 10X Chromium platform were processed using the standard CellRanger pipeline (version 5.0.0). Reads were aligned to GRCm38/mm10 reference. Cell-gene count matrices were analyzed using a combination of published packages and custom scripts centered around the scanpy/AnnData ecosystem ^85^. scRNA-seq data from eIF2αS/S and eIF2αA/A mouse lung tumor were dehashed and compiled into a combined count matrix. Cells with less than 500 Unique molecular identifiers (UMIs), more than 20% mitochondrial UMIs, and low complexity based on the number of detected genes vs number of UMIs were removed. Doublets were filtered using DoubletFinder^86^. Highly variable features were selected using a variance stabilizing transformation, and dimensionality reduction was performed on normalized, log2-transformed count data using principal component analysis. The dimensionality-reduced count matrices were used as input for UMAP-embedding and unsupervised clustering with the Leiden algorithm; bbknn was used to control for batch effects ^87^. To ensure that treatment does not bias cell state assignment, unsupervised clustering was performed only on control cells (eIF2αS/S or eIF2αS/S vehicle-treated) and used as input to train a logistic regression (logit) classifier which was used to assign each cell to a cell state. This approach ensured that the cell states defined in the study are representative of states previously identified in unperturbed mouse KP lung tumors ^23^. Normalized expression data were first MaxAbs scaled to give each gene equal weight. Gene signature scores were then calculated using the score genes function in Scanpy. The normal cell signatures were from MSigDB ^47,83^, and KP tumor cell signatures were from Ref. ^23^. Bhattacharyya distances were calculated using the bhattacharyya function from the python package distances 1.5.6).

### Analysis of gene expression along the LUAD evolution trajectory

We implemented CellRank ^40^ to uncover the dynamic cell state transitions from single cell data. CellRank uses a generalized additive model (GAM) to smooth expression along the AT2-like to Pre-EMT or Mito-Dysfunction trajectory. Imputed expressions were used to generate this visual, and expression was normalized to a range of [0,1]. The order of genes was determined by the expression peak along the terminal state probability continuum. Gene trend curves were generated using the built-in plotting method provided by CellRank.

### Transcription factor regulatory module (regulon) analysis

To detect regulatory programs differentially active across cell populations, we implemented the SCENIC ^48^ workflow for detecting activating transcription factor regulatory modules (regulons) with default parameters and provided reference databases (mm10 transcription factor targets and 500bp-up-100bp-down gene-motif rankings). In addition to the standard SCENIC approach to detecting condition-specific regulons in discrete comparisons, we also detected top regulons associated with the cluster using the Pearson correlations.

### Analysis of public datasets

Processed scRNA-Seq profiles from human LUAD tumors were downloaded from the CZI cellxgene portal ^58^. Our analysis was restricted to LUAD samples, focusing only on cells labeled by the original study authors as cancer cells. For murine data, we obtained processed scRNA-Seq profiles from mouse lung samples from GSE122332 ^88^. In this dataset, our analysis was limited to alveolar type II (AT2) and alveolar type I (AT1) cells, as annotated by the study authors.

### Human LUAD survival analysis

TempO-seq targeted sequencing-based RNA expression sequencing was performed on 706 non-mucinous LUAD cores from 23 FFPE TMA slides. Samples were scraped and lysed using the TempO-Seq 2X Lysis Buffer. Sequencing was performed based on the Human Whole Transcriptome v2.1 panel with standard attenuators. Tempo-Seq-RTM software package was used to pre-process data, reads passing QC were aligned using the STAR algorithm ^89^ to a pseudo-transcriptome corresponding to the gene panel used. Gene isoforms were merged, and genes detected in less than a third of samples were removed before normalizing using upper-quartile method and a log2 transformation. Gene signature expression levels were calculated using single-sample GSEA method as implemented in the GSVA R package (v1.51-17) ^90^based on the genes with FDR<0.05 and logFC>1.5. Cox proportional hazard regression was used to perform survival analysis as implemented in survival R package (v3.6-4). Driver mutation information was available for 469 patients of which 180 had a driver KRAS mutation. Samples were stratified between high and low p-eIF2α using median H-score as a cut-off.

### Duplex immunohistochemistry for p-eIF2α and Cytokeratin AE1/AE3

The expression of p-eIF2α and cytokeratin AE1/AE3 was assessed using an optimized duplex immunohistochemistry protocol performed on a Ventana DISCOVERY ULTRA automated staining platform (Roche Tissue Diagnostics). This assay was validated on a comprehensive tissue microarray (TMA) of human lung adenocarcinoma. High-resolution digital imaging was conducted using a Hamamatsu NanoZoomer system.

Antigen retrieval was performed under high pH conditions (pH 9.0, Roche Tissue Diagnostics, 06414575001) for 64 minutes at 95°C to unmask epitopes. The primary antibody against p-eIF2α (Cell Signaling Technology, 3398) was applied at a dilution of 1:25 and incubated for six hours at room temperature. Detection was achieved using UltraMap anti-rabbit HRP (Roche Tissue Diagnostics, 05269717001) for 28 minutes at 37°C, and visualization was accomplished with DAB chromogen (Roche Tissue Diagnostics, 05266645001) for 8 minutes at 37°C. Signal enhancement was performed using the DISCOVERY Amp TSA HQ Kit (Roche Tissue Diagnostics, 06472320001), incubated for 12 minutes at 37°C followed by Discovery Amplification anti-HQ HRP multimer (Roche Tissue Diagnostics, 06442544001) for 12 minutes. To allow for sequential staining, antibody denaturation was conducted in a pH 6.0 buffer (Roche Tissue Diagnostics, 05424542001) for 8 minutes at 100°C. Slides were treated with DISCOVERY inhibitor (Roche Tissue Diagnostics, 07017944001) for 20 minutes before proceeding to the second round of staining. The primary antibody for cytokeratin AE1/AE3 (Leica Biosystems, NCL-L-AE1/AE3) was applied at a dilution of 1:250 and incubated for 28 minutes at 37°C. Detection was performed using OmniMap anti-mouse HRP (Roche Tissue Diagnostics, 05269652001) for 12 minutes at 37°C, followed by development with a purple chromogen (Roche Tissue Diagnostics, 07053983001) for 16 minutes at 37°C. A haematoxylin counterstain (Roche Tissue Diagnostics, 05277965001) was applied for 8 minutes to provide nuclear contrast.

Following staining, the slides were digitized using a Hamamatsu NanoZoomer whole-slide scanner using a 40x objective. Digital images were analyzed using Visiopharm. A tissue segmentation classifier was generated using a Threshold module to segment tissue into tumor and stroma regions of interest. Cells were segmented using the Cell Classification module, and an H-Score was generated by manually assigning intensity cut offs. H-Score thresholds were verified by a pathologist.

## ACKNOWLEDGMENTS

We acknowledge the excellent expertise of Chris Young and Naciba Benlimame at the core facility of the *Lady Davis* Institute and Segal Cancer Centre, Jewish General Hospital, for their assistance with flow cytometry and immunohistochemistry services, respectively. This work was funded by the Canadian Institutes of Health Research (CIHR) through a grant awarded to A.E.K. (PJT-168864), National Institute of Health (NIH) DK53307 to MH, NIH R01-CA270116 and Josie Robertson and Rita Allen Scholarships to TT, NIH 2P01-CA165997 1 R01 CA268597 to CK, and Mazumdar-Shaw Chair endowment (University of Glasgow) to JLQ. JYZ is supported by a studentship award from *Le Fonds de recherche du Québec* (*FRQS*), and HK is a recipient of the Faculty of Medicine award from McGill University.

## AUTHOR CONTRIBUTIONS

Conceptualization: AEK and TT. Methodology: SD, JYZ, SW, NG, HK, JEC, NP. Software: SD, JEC, NP Validation: CK, MH, PW, NS, JLQ, Formal Analysis: SD, JYZ, JEC, NP Investigation: SD, JYZ, SW, NG, HK, JEC, NP. Data Curation: SD, JEC, NP. Writing: AEK, SD, JYZ, MH, PW, NS, JLQ, TT. Visualization: SD, JYZ, NG, NP. Supervision: AEK, TT, NS, CK. Project Administration: AEK

## DATA AVAILABILITY

For data availability, the dataset can be found on GEO under accession number GSE287513.

---

**Extended Data Fig. 1.**
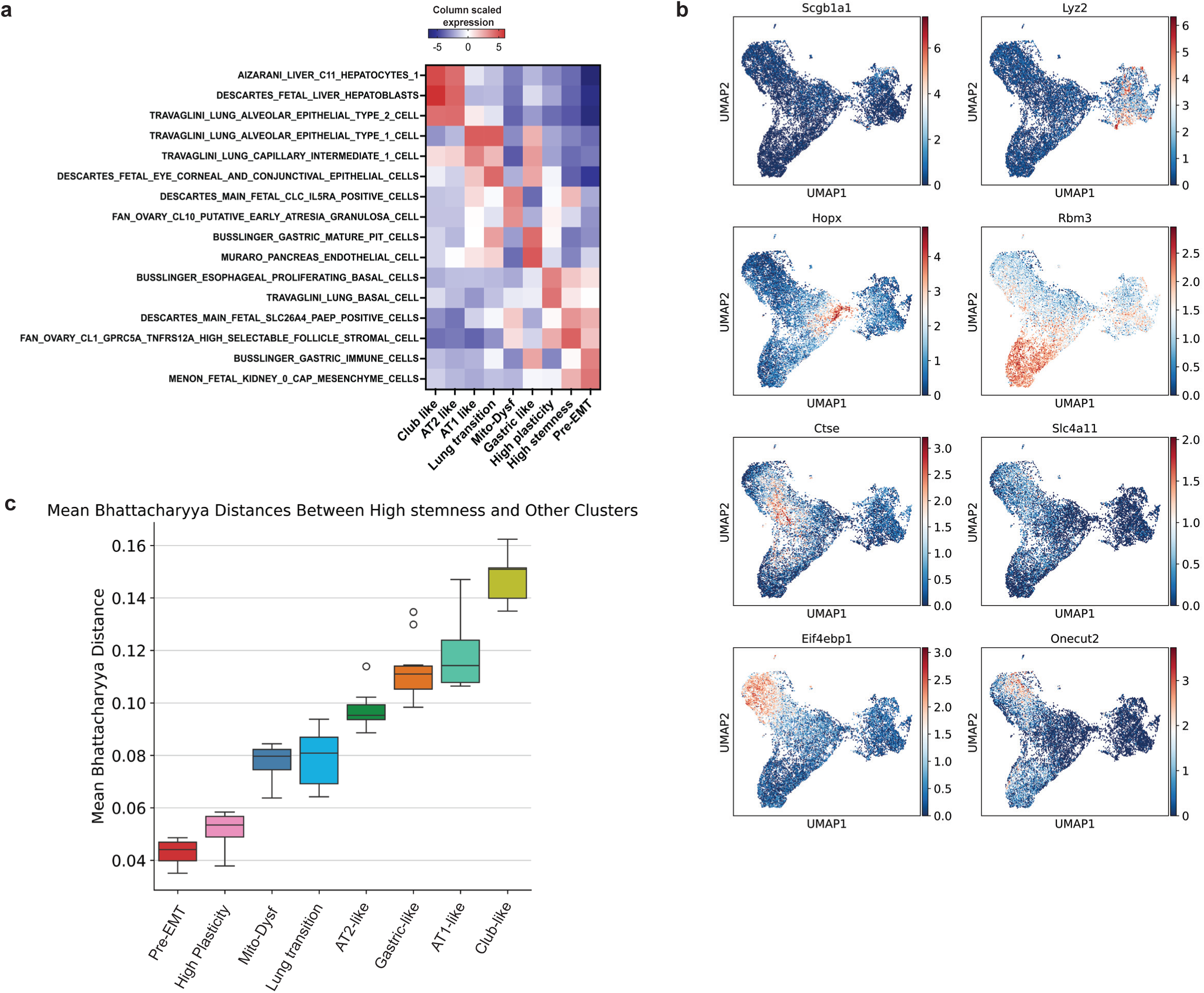
Characterization of cluster cells in eIF2α^S/S^ and eIF2α^A/A^ KP tumors. (**a**) Heatmap displaying scaled feature scores of MSigDB canonical cellular features across annotated cell types, highlighting distinct patterns of feature enrichment in specific cell types. These cell types are organized sequentially according to their roles in tumor development. (**b**) Gene markers were identified for each Leiden cluster in the processed scRNA-seq latent space. The mean expression level of these marker genes is represented by color. (**c**) The Bhattacharyya distance quantifies the similarity between the high-stemness other inferred cell lineages, revealing the molecular and phenotypic relationships of the different cluster and their contributions to tumor heterogeneity and evolution.

**Extended Data Fig. 2.**
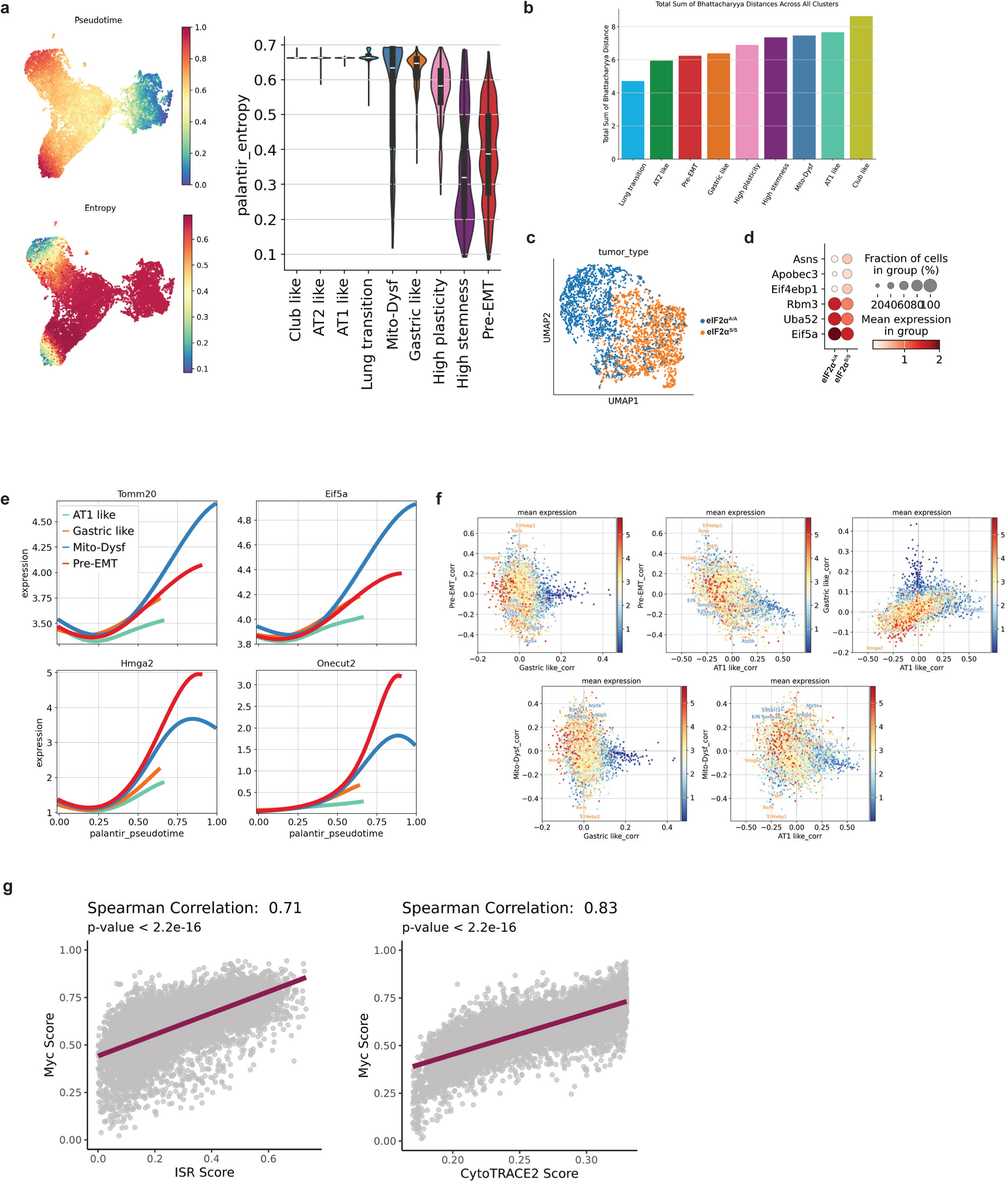
Critical turning points in the evolution of eIF2α^S/S^ and eIF2α^A/A^ tumor cells. (**a**) CellRank analysis for the identification of the entropy of fate probability as a measure of the uncertainty in cell fate decisions. Entropy is maximal when there is no fate bias and decreases as cells move toward terminal states. This analysis highlights the dynamics of cell fate decisions and their contributions to tumor evolution and heterogeneity. The accompanying violin plot illustrates the distribution of entropy across different cell clusters. (**b**) The total Bhattacharyya distance quantifies the overall similarity between a selected cluster and all other inferred cell lineages. (**c**) The UMAP plot visualizes the lung transition cluster derived from eIF2α^S/S^ and eIF2α^A/A^ tumors, encompassing a total of 3,909 cells. (**d**) The dot plot of top DEGs in eIF2α^S/S^ and eIF2α^A/A^ tumors from panel C. (**e**) The smoothed gene expression trends of *Tomm20, Eif5a, Onecut2* and *Hmga2* along pseudotime are visualized, with each trend weighted by CellRank fate probabilities. The data shows the dynamic changes in gene expression over time, highlighting the correlation of these genes with predicted cell fate trajectories. *Tomm20, Eif5a* are key driver genes for Mito-Dysf cluster, while *Onecut2* and *Hmga2* are the key driver genes for pre-EMT cluster. (**f**) The scatter plot illustrates the correlation of each gene with different fates, plotted along the x-axis and y-axis. Each point represents a gene, colored according to its average expression across all cells. (g) Correlation analyses between Cytotrace stemness scores, ISR activity, and MYC activity. Spearman’s correlation test was used to calculate p-values.

**Extended Data Fig. 3.**
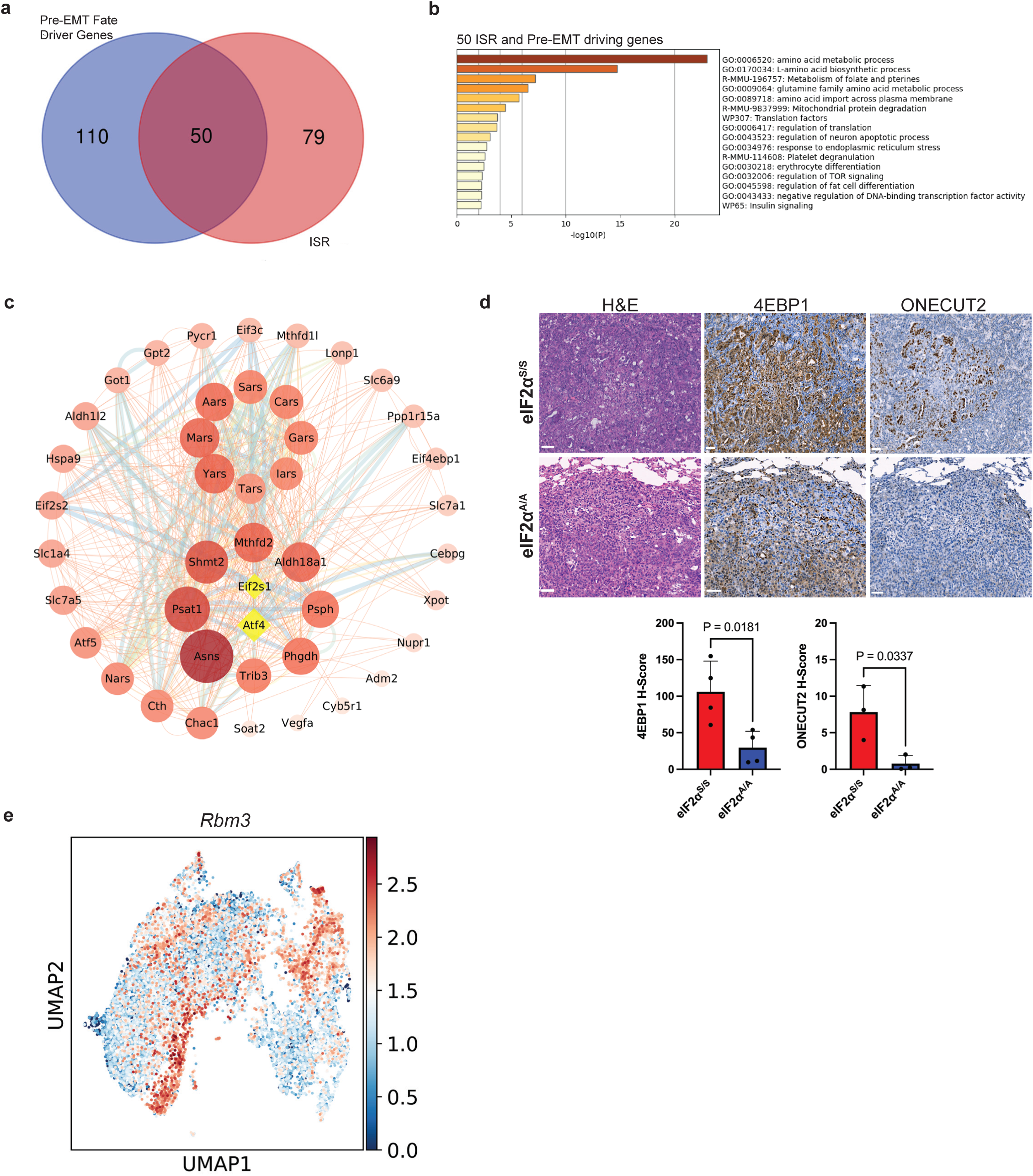
The ISR signaling plays a critical role in driving the evolution of AT2-like into pre-EMT states. (**a**) The Venn diagram illustrates the percentages of unique and overlapping gene sets between the top pre-EMT fate driver genes identified by CellRank and the ISR gene set. (**b**) The pathway enrichment analysis of the 50 common genes identified in panel A highlights key biological pathways and processes enriched in this gene set. The analysis provides insights into the shared mechanisms linking ISR signaling and pre-EMT fate-driving genes. (**c**) The Protein-Protein Interaction (PPI) plot illustrates the network of the 50 common genes identified in panel A, with interaction strength represented by color intensity and circle size. Nodes with higher interaction degrees are highlighted with larger circles and more intense colors, indicating their central roles in the network. *Atf4* and *Eif2s1* nodes are manually integrated into the plot, emphasizing their critical regulatory roles and connections within the network. (**d**) The IHC staining analysis of the high-stemness/pre-EMT marker 4EBP1 (*n* = 4 mice) and the pre-EMT marker ONECUT2 (*n* = 3 mice) was conducted in in eIF2α^S/S^ and eIF2α^A/A^ mouse tumors; scalebars, 50 μm. Statistical comparisons between the groups were performed using a two-tailed unpaired t-test, with results presented as mean ± SD. (**e**) UMAP analysis merging cells from eIF2α^S/S^ and eIF2α^S/A^ genotypes, with each dot representing a single cell. Cells are colored by *Rbm3* expression levels, ranging from low (blue) to high (red).

**Extended Data Fig. 4.**
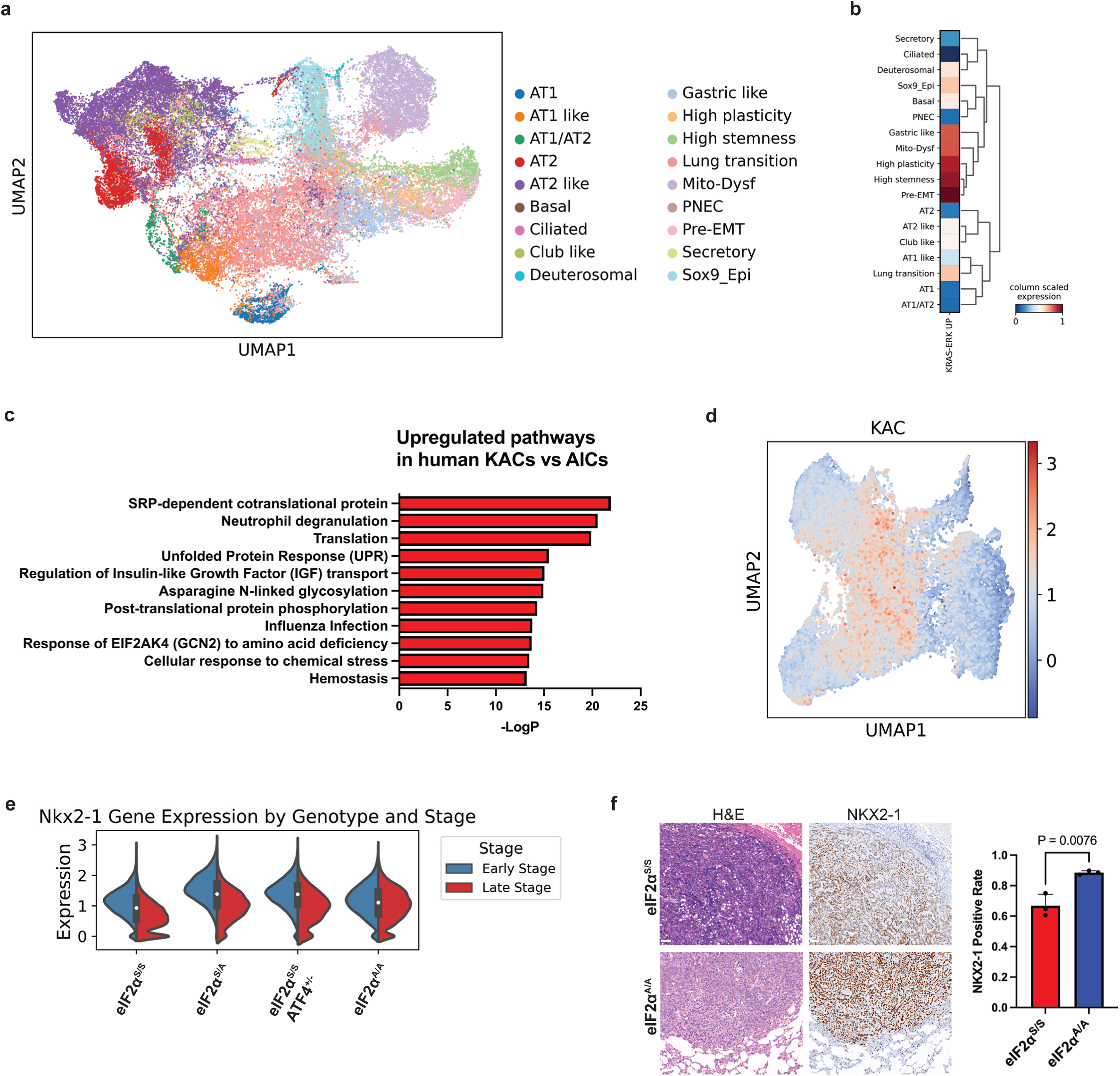
The lung transition cluster serves as a critical node in the ISR-driven evolution of AT2-like cells into dedifferentiated tumor states. (**a**) The UMAP plot illustrates the integration of tumor samples with normal lung atlas datasets, providing a comparative view of cellular populations. The integration reveals tumor-specific cellular states, lineage transitions, and deviations from normal lung tissue, aiding the analysis of tumor heterogeneity (n=65,357 cells). (**b**) Heatmap shows the KRAS-ERK dependent genes expression in each cell cluster. (**c**) GSEA comparing KRT8+ alveolar intermediate cells (KACs) and alveolar intermediate cells (AICs) from the GSE222901 dataset identifies pathways and biological processes enriched in each cell type, including ISR-mediated pathways. (**d**) The UMAP plot visualizes all datasets obtained from KP eIF2α^S/S^, eIF2α^A/A^, eIF2α^S/A^ and eIF2α^S/S^ ATF4^+/-^ tumors, with cells colored based on the expression levels of KAC marker genes. The gradient reflects the intensity of marker gene expression, highlighting regions enriched with KACs within the tumor microenvironment. (**e**) Violin plot shows *Nkx2-1* expression in ISR tumor datasets, categorized by early and late-stage tumors. (**f**) IHC staining of NKX2-1 in tumors from eIF2α^S/S^ and eIF2α^A/A^ mice (n = 3 mice per genotype); scalebars, 50 μm. Statistical analysis was performed using a two-tailed unpaired t-test, and data are presented as mean ± SD

**Extended Data Fig. 5.**
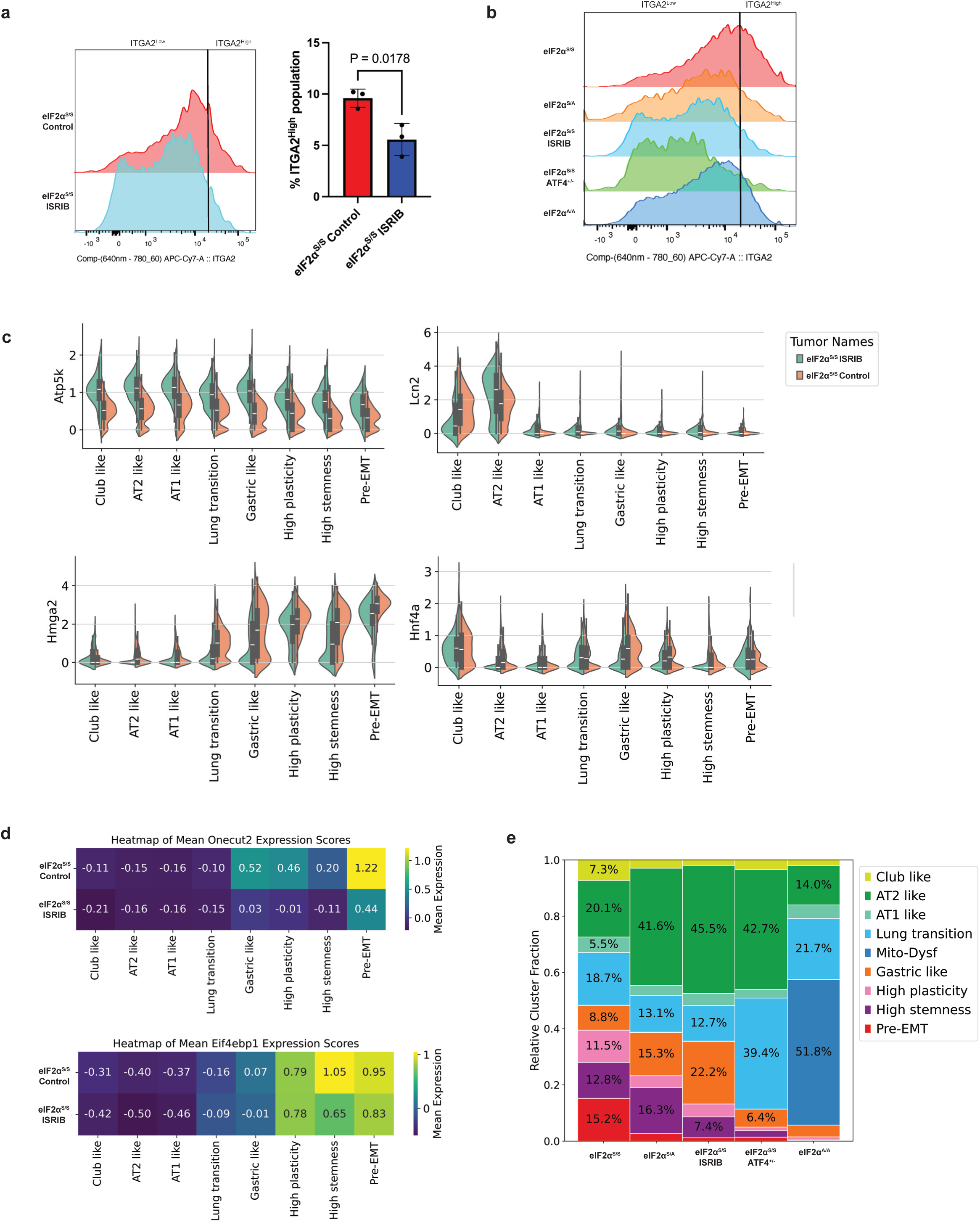
ISRIB treatment enhances mitochondrial and AT2 markers while reducing dedifferentiated tumor markers. (**a**) Representative histograms depicting the distribution of ITGA2 expression in vehicle-treated and ISRIB-treated eIF2α^S/S^ tumors. Statistical analysis was conducted using a two-tailed unpaired t-test (n = 3 mice per group) (**b**) Detection of ITGA2 expression by flow cytometry in primary tumor populations of the indicated genotypes. (**c**) Violin plots illustrating the expression of *Atp5k*, *Lcn2*, *Hmga2*, *Hnf4a* marker genes in tumors treated with either ISRIB or vehicle control. (**d**) Heatmap showing the mean expression levels of *Onecut2* and *Eif4ebp1* across clusters in tumors treated with either vehicle or ISRIB. (**e**) Cluster cell proportions in LUAD tumors of the indicated genotypes visualized after integrating all ISR sequencing datasets.

**Extended Data Fig. 6.**
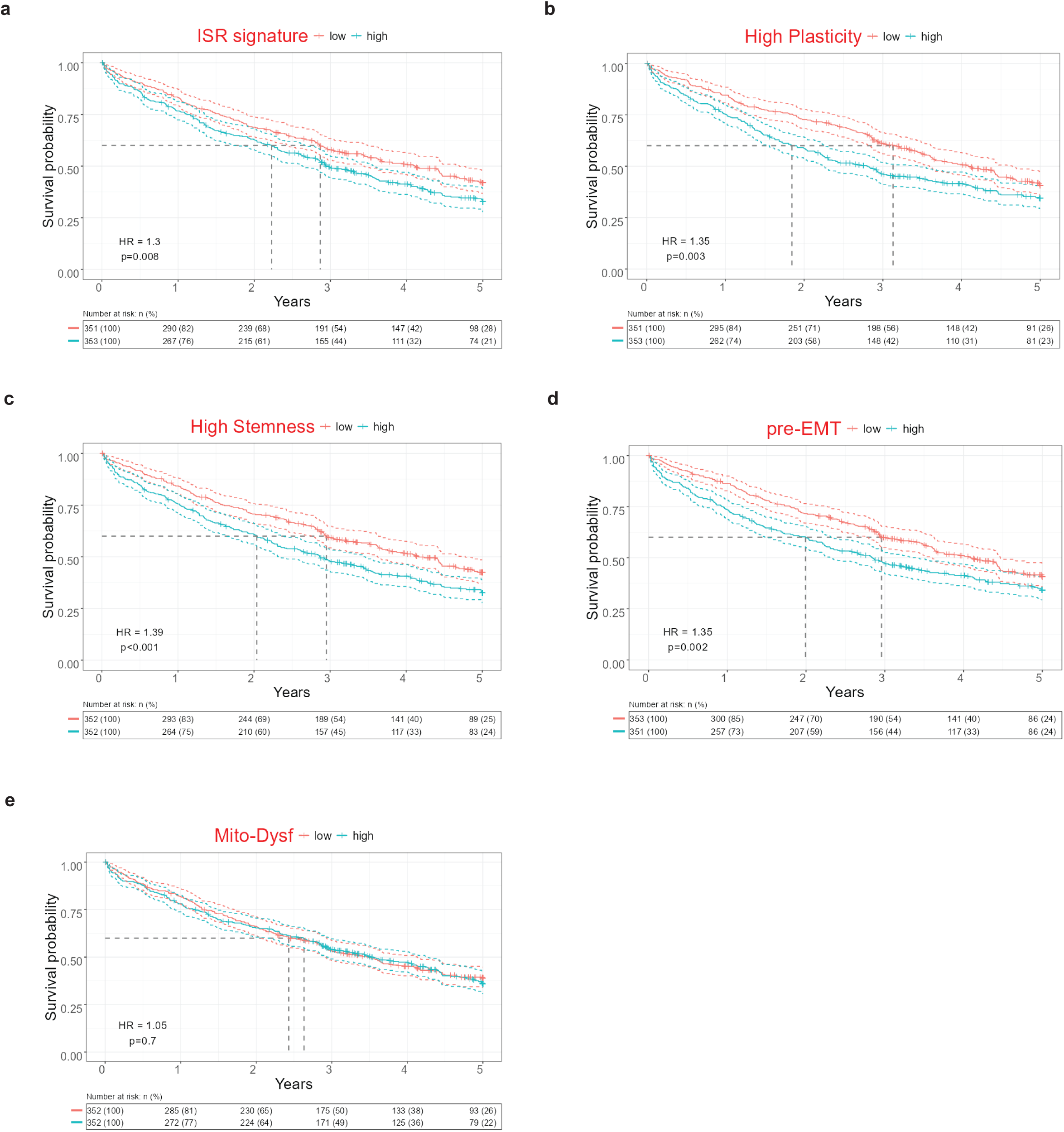
Marker genes of ISR-driven dedifferentiated clusters are identified in LUAD patient samples and associated with poor survival outcomes. **(a-e)** Kaplan-Meier plots illustrating the survival impact of cluster marker gene expression in patient samples from a cohort of 706 non-mucinous LUAD cases. Analysis reveals that elevated expression levels of the ISR signature (**a**), high plasticity (**b**), high stemness (**c**), and pre-EMT programs (**d**) are significantly associated with poorer survival outcomes. In contrast, the Mito-Dysf expression program shows no significant association with survival (**e**).

**Extended Data Fig. 7.**
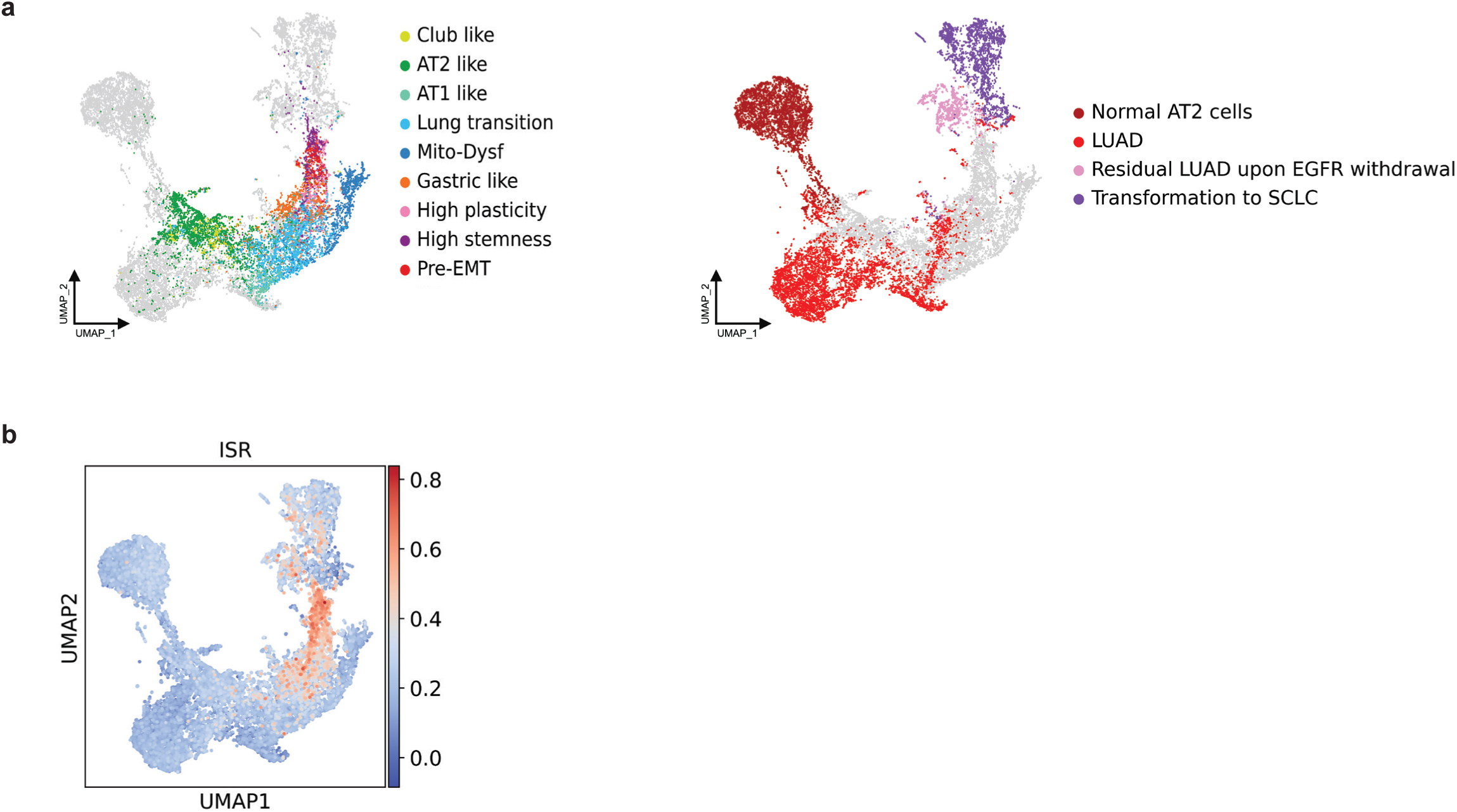
UMAP Analysis of Combined ISR-KP and HT Mouse Model Datasets. (**a**) UMAP merging the datasets of the ISR-KP model from this study with the histological transformation (HT) mouse model of EGFR-driven LUAD Ref ^25^. Colors represent annotated cell types identified in this study (left panel) and in EGFR-driven LUAD tumors of the HT model (right panel). AT2-like cells in the ISR-KP model show similarities to normal AT2 cells and LUAD cluster cells in the EGFR-driven LUAD model. The high-stemness cell cluster in the ISR-KP model (left panel, purple) corresponds to cells located above the residual LUAD cluster cells, exhibiting characteristics of undifferentiated cells. These also align with SCLC-transformed cells in the EGFR-driven cancer (right panel). The combined dataset includes a total of 25,828 cells from both models. (**b**) UMAP showing that expression of ISR dependent genes is increased in high-stemness cluster and residual LUAD cells of the EGFR-driven LUAD model.

